# Whole-brain clearing reveals region- and cell type-specific imbalances in inhibitory neurons in a mouse model for Kleefstra Syndrome

**DOI:** 10.1101/2025.09.15.676416

**Authors:** Moritz Negwer, Ilse van der Werf, Astrid Oudakker, Hans van Bokhoven, Dirk Schubert, Nael Nadif Kasri

## Abstract

GABAergic inhibition is essential for balanced brain function and is frequently disrupted in neurodevelopmental disorders such as autism spectrum disorder (ASD). The inhibition is generated by a diverse population of GABAergic interneurons, whose composition and density differs throughout the brain. Here, we applied an unbiased whole-brain clearing and light-sheet imaging approach to systematically map the distribution of the three major GABAergic neuron classes - Parvalbumin-positive (PV⁺), Somatostatin-positive (SST⁺), and Vasoactive Intestinal Polypeptide-positive (VIP⁺) cells - across the mouse brain in a model of Kleefstra Syndrome (*Ehmt1^+/−^*), a monogenic intellectual disability disorder with a strong ASD component. Analyzing 895 brain regions we identified widespread, cell type- and region-specific alterations in GABAergic classes. Notably, we observed increased VIP⁺ neuron density in the *Ehmt1^+/−^* cortex and decreased SST⁺ neuron density in sensory cortical and subcortical regions. In the basolateral amygdala (BLA), PV⁺ neurons exhibited precocious maturation already at the juvenile stage, which persisted into adulthood and was associated with enhanced inhibitory input onto BLA principal neurons. We here demonstrate that *Ehmt1* haploinsufficiency results in region- and cell-type specific changes throughout the brain. These results underscore the value of whole-brain, high-resolution mapping approaches in uncovering previously unrecognized patterns of neural vulnerability in neurodevelopmental disorders.

## Introduction

Brain function relies on a precise balance between excitation and inhibition. In mice, inhibition is provided by approximately 16% of neurons that produce the neurotransmitter gamma-aminobutyric acid (GABA; Bakken et al., 2021; Fishell and Kepecs, 2020; Gouwens et al., 2020, 2019; Huang and Paul, 2019; Yao et al., 2021). The percentage of GABAergic neurons is higher in humans than in mice at ca. 33% (Bakken et al., 2021), however with generally conserved cell classes (Lee et al., 2023) and a conserved excitatory/inhibitory (E/I) balance (Loomba et al., 2022). GABAergic neurons exhibit remarkable diversity, which at the highest level of detail can be distinguished into up to 80 subclasses (Gouwens et al., 2020). A widely accepted classification groups these neurons into three main classes - Parvalbumin-expressing (PV^+^), Somatostatin-expressing (SST^+^), and Vasoactive Intestinal Polypeptide-expressing (VIP^+^) neurons - which together account for about 80% of cortical GABAergic neurons (Bakken et al., 2021; Gouwens et al., 2020; Rudy et al., 2011; Tremblay et al., 2016). These classes play distinct roles in the canonical cortical microcircuits (Favuzzi et al., 2019; Fisher et al., 2024; Piet et al., 2024): PV^+^ neurons provide fast, perisomatic inhibition to pyramidal neurons (Klausberger and Somogyi, 2008), SST^+^ neurons (e.g. cortical Martinotti cells) target dendrites to regulate input (Kawaguchi and Kondo, 2002), and VIP^+^ neurons preferentially inhibit other GABAergic neurons, disinhibiting pyramidal cells (Rachel et al., 2025). Similar populations of GABAergic neurons exist in the cortical areas of the amygdala, with additional striatal-like populations in its medial compartments (Hájos, 2021; Hochgerner et al., 2023). The identity of the GABAergic neuron classes is generated and maintained by epigenetic regulation (Achim, 2024; Fishell and Kepecs, 2020; Li et al., 2026; Rhodes et al., 2022). Therefore, GABAergic neurons are vulnerable to disruption in epigenetic regulator genes (Mossink et al., 2021), which feature prominently in neurodevelopmental disorders (Marín, 2024; Tang et al., 2021).

One such epigenetic factor, Euchromatic Histone Methyltransferase 1 (*EHMT1*), plays a critical role in neurodevelopment. Haploinsufficiency of the *EHMT1* gene causes Kleefstra syndrome (KLEFS1, MIM: 610253; Rots et al., 2024), a rare neurodevelopmental disorder characterized by intellectual disability, autistic traits, anxiety, and sleep disruptions (Kleefstra, 2005; Kleefstra et al., 2006; Vermeulen et al., 2017b; Willemsen et al., 2011). The EHMT1 protein functions as part of the NRSF/REST complex to regulate gene expression through mono- and dimethylation of histone H3 at lysine 9 (H3K9me1/2; (Tachibana et al., 2008, 2005, 2002). In mice, *Ehmt1* haploinsufficiency leads to a range of neurological and molecular abnormalities, including overactive glutamatergic signaling (Balemans et al., 2014, 2013, 2010; Benevento et al., 2017, 2016, 2015; Schut et al., 2020), neuroinflammation (Yamada et al., 2021), and dysregulated expression of genes dependent on three-dimensional chromatin structures, such as clustered protocadherins (Iacono et al., 2018).

We previously demonstrated that PV^+^ neurons in the sensory cortices of *Ehmt1*^+/−^ mice mature more slowly than in wild-type mice, but these studies were limited to one GABAergic class in three main sensory cortical areas, layers 2/3 and 4 (Negwer et al., 2020). To comprehensively address how *Ehmt1* haploinsufficiency affects GABAergic neurons, we conducted a brain-wide analysis of the three major GABAergic classes, i.e. PV^+^, SST^+^ & VIP^+^ We stained, cleared and imaged whole mouse brain hemispheres following the iDISCO+ protocol (Kirst et al., 2020; Renier et al., 2016, 2014) and used FriendlyClearmap for analysis (Negwer et al., 2022). We observed substantial class-specific differences in GABAergic neuron densities in *Ehmt1*^+/−^ brains, including an unexpected early increase in PV^+^ neuron density in the basolateral amygdala (BLA) as well as a general reduction in SST^+^ density and higher VIP^+^ density in the cortex. We confirmed that the higher PV^+^ neuron density in the *Ehmt1^+/−^* amygdala leads to higher inhibitory drive already at P14 and persists into adulthood. However, this does not translate to obvious behavioral changes following a social novelty challenge.

## Results

### Region-specific cell density differences for all three main GABAergic neuron types in *Ehmt1*^+/−^ mice

We chose an unbiased, brain-wide approach to quantify the density of the three major GABAergic neuron classes in the brain (Fig. 1). To this end, we crossed *Ehmt1*^+/−^ mice with tdTomato reporter lines for the Parvalbumin-expressing (PV^+^), Somatostatin-expressing (SST^+^), and VIP-expressing (VIP^+^) classes of GABAergic neurons (Fig. 1a). We specifically chose a reporter mouse strain because this allowed us to visualize several GABAergic neuron classes with a single immunostaining with high signal/noise ratio. We then harvested brains of *tdTom^+^*/*Ehmt1^+/−^* and *tdTom^+^*/*Ehmt1^+/+^* littermates. We selected the timepoints according to the developmental trajectory of the respective cell type: Adolescence (P28) for VIP^+^ and SST^+^, when those cells are developmentally mature. PV^+^ neurons have an especially long postnatal developmental trajectory, therefore we picked an early timepoint (P14) when PV is just starting to be expressed in the cortex, and an adult timepoint (P56) when PV^+^ cells are mature (Negwer et al., 2020). We then used the iDISCO+ staining/clearing protocol (Kirst et al., 2020; Renier et al., 2016, 2014) with slight modifications (Negwer et al. 2022; Fig. 1b), which ensured complete and even anti-tdTomato staining throughout the hemisphere (supplementary Fig. 1). The resulting image stacks were then processed with FriendlyClearMap (Negwer et al. 2022; Fig 1c) to generate brain-wide density maps for PV^+^, SST^+^ and VIP^+^ neurons. With FriendlyClearMap, we mapped our data to the Allen Brain Atlas CCF3 (Wang et al., 2020) for the adult mice, and to the developmental brain atlas (Newmaster et al., 2020) for the P14 PV^+^ and P28 SST^+^ and VIP^+^ brains, to get GABAergic neuron densities for 895 brain regions in the forebrain and midbrain.

**Figure 1:**
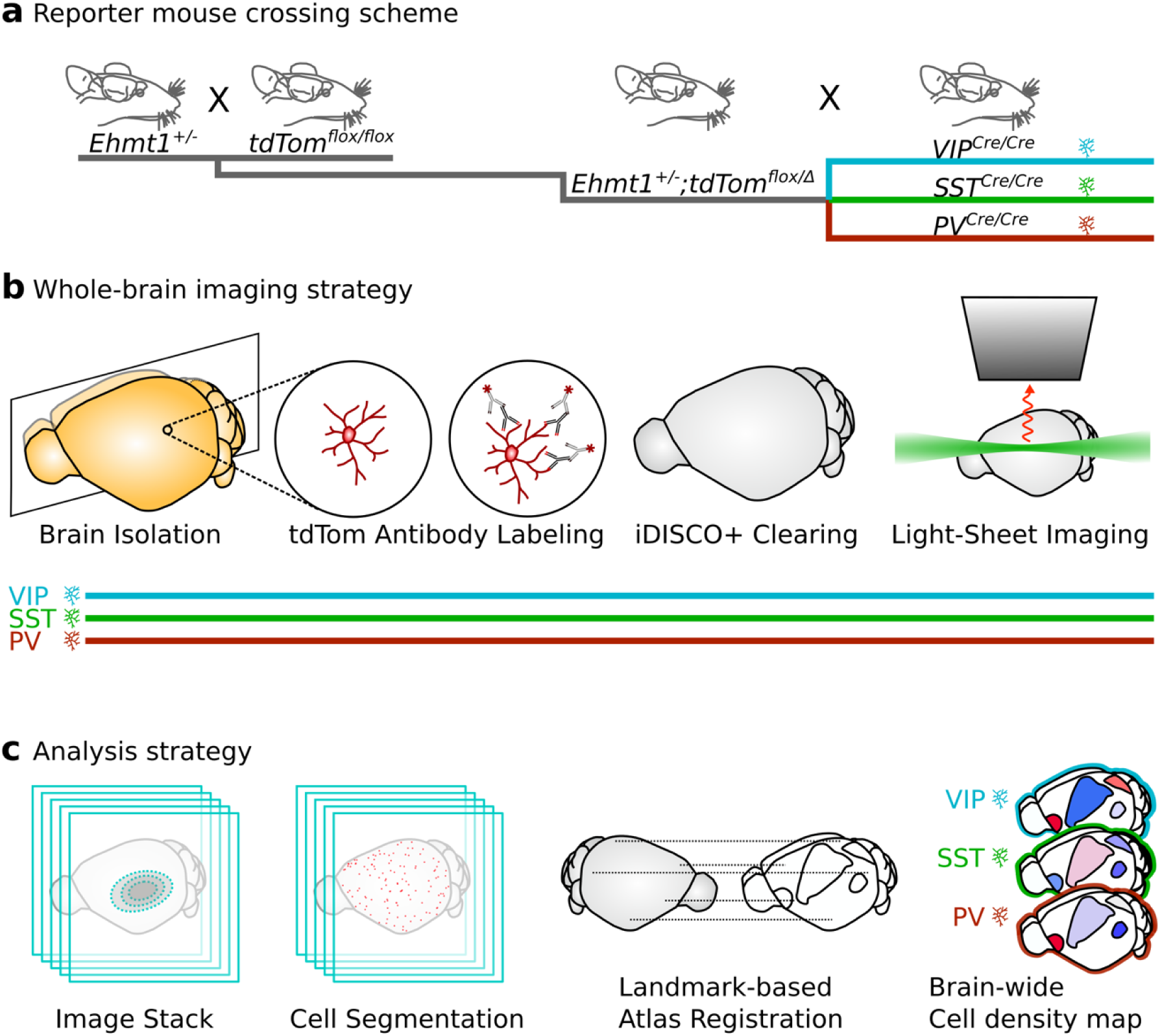
Experimental Strategy. **a)**, we generated reporter lines for the three main GABAergic neuron types: Vasoactive Intestinal Polypeptide-positive (VIP^+^), Somatostatin-positive (SST^+^), and Parvalbumin-positive (PV^+^). We crossed *Ehmt1^+/−^* mice with *tdTomato^flox^* reporter mice (for two generations), and used the *Ehmt1^+/−^;tdTom^flox/flox^* offspring for a second crossing with Cre driver lines: *Vip^tm1(cre)Zjh/J^* (*VIP^Cre/Cre^*), *Sst^tm2.1(cre)Zjh/J^* (*SST^Cre/Cre^*), or *B6.129P2-Pvalb^tm1(cre)Arbr/J^* (*PV^Cre/Cre^*). The offspring expressed tdTomato in either VIP, SST or PV neurons, and we used *Ehmt1^+/+^* and *Ehmt1^+/−^* as control and mutant animals, respectively. **b)**, we then sacrificed the animals, removed the brain as hemispheres, and used a modified iDISCO technique to label tdTomato with antibodies tagged with Alexa647 fluorophores, followed by Methanol/DCM/DBE clearing. We then imaged the cleared hemispheres with a Miltenyi Ultramicroscope II at 2.2x for autofluorescence (ex. 488nm) and tdTomato (ex. 640nm). **c)** We then processed the image stacks with FriendlyClearMap (Negwer et al., 2022), using Ilastik for cell segmentation and a manual landmark-based atlas registration. With this approach, we generated whole-brain cell density maps for VIP^+^, SST^+^ and PV^+^ neurons.

VIP^+^ neurons comprise approximately 12% of all cortical GABAergic neurons (Gouwens et al., 2020, 2019; Tremblay et al., 2016). VIP^+^ neurons appear mostly in the cortex, with the large majority in layers 2/3 (Prönneke et al., 2015). They are marked by the neuromodulatory peptide Vasoactive Intestinal Polypeptide (VIP) which they co-release with GABA (Apicella and Marchionni, 2022). In the canonical cortical circuit, their function is thought to be disinhibition of other GABAergic neurons, notably SST^+^ (Piet et al., 2024; Williams and Holtmaat, 2019). With our brain-wide iDISCO+ screening, we indeed found that VIP^+^ neurons in mouse brains at P28 were mostly localized in the cortex in both wild-type and *Ehmt1^+/−^* mice (Fig. 2a). However, we did find a strikingly higher VIP^+^ neuron density in the *Ehmt1^+/−^* visual, somatosensory and auditory cortices (Range: 126-195% of *Ehmt1^+/+^*; Fig. 2a-c, Fig. 3a; supplementary Fig. 1, supplementary table 1), where VIP^+^ density is comparatively low in wild-type mice (Kim et al., 2017; Negwer et al., 2022). We found that the increase in sensory areas was roughly evenly split between supragranular (layers 2/3 and 4) and infragranular (layers 5 and 6) VIP^+^ neurons (Fig. 2C, Fig. 3b, supplementary table 1).

**Figure 2:**
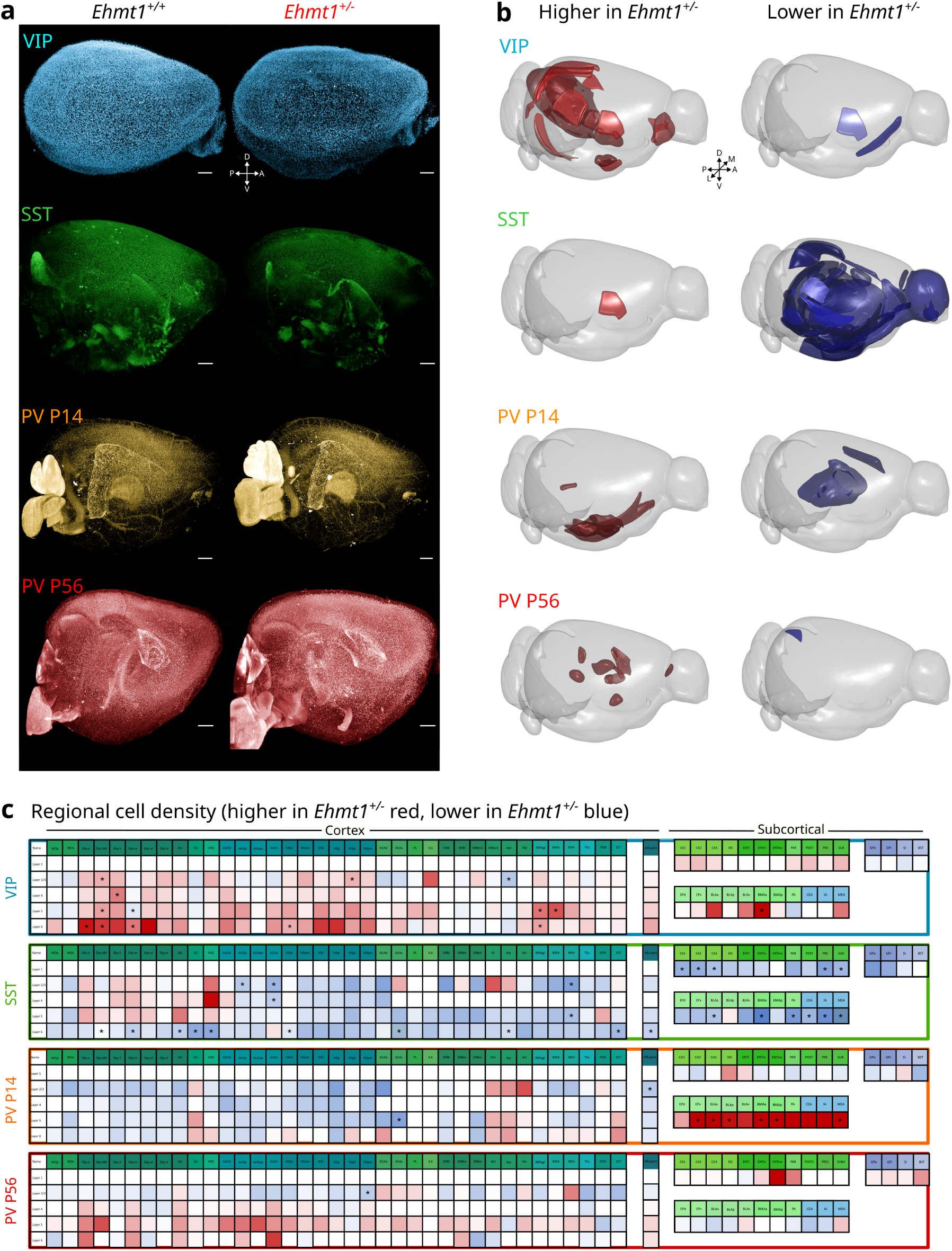
GABAergic neuron distribution differences in the *Ehmt1^+/−^* mouse brain found with iDISCO and FriendlyClearMap. **a)** Representative whole-hemisphere 3D images of the three main GABAergic neuron types in *Ehmt1^+/+^* (left) and *Ehmt1^+/−^* (right) brains. **b)** Significantly different areas where *Ehmt1^+/−^* mice have higher (red) or lower (blue) cell densities compared to *Ehmt1^+/+^*. We find higher VIP^+^ density in *Ehmt1^+/−^* retrosplenial, somatosensory, auditory and entorhinal cortices. We find significantly lower densities in the insular cortex. See supplementary table 1 for details. We find reduced SST^+^ density in *Ehmt1^+/−^* amygdala, thalamus (several nuclei each), hippocampus, auditory and somatosensory cortices. See supplementary table 2 for details. At P14, we find lower PV^+^ density in *Ehmt1^+/−^* in the anterior cingulate cortex, but surprisingly higher in several nuclei of the amygdala. See supplementary table 3 for details. In adulthood (P56), we find significantly reduced PV^+^ density in the *Ehmt1^+/−^* Posteromedial Visual cortex (L2/3), but higher PV^+^ density in *Ehmt1^+/−^* in subcortical nuclei in the commissure, pretectum, and Hypothalamus. Please see supplementary table 4 for details. Scale bars = 500 µm. A/P = Anterior/Posterior, D/V = Dorsal/Ventral, M/L = Medial/Lateral. **c)** Density of tdTom-tagged GABAergic neurons in *Ehmt1^+/−^* as percentage of *Ehmt1^+/+^* in cortical regions and selected amygdalar and hippocampal fields. Blue indicates that *Ehmt1^+/−^*density is lower compared to *Ehmt1^+/+^*, and red indicates a higher density. Asterisks indicate significant differences in nonparametric Mann-Whitney U tests. *Ehmt1^+/−^* has higher VIP^+^ densities in the sensory cortical areas (especially somatosensory) and in the retrosplenial area, but also in the basolateral and basomedial amygdala. The layer overview indicates a shift towards the infragranular layers in *Ehmt1^+/−^*. SST^+^ neuron density is lower in most of the *Ehmt1^+/−^* cortex (with the exception of the somatosensory areas), particularly in layer 6. SST^+^ density is also lower in the *Ehmt1^+/−^* BLAa, BMAa, CEA and IA amygdalar regions. PV^+^ density in the *Ehmt1^+/−^* at P14 is similarly lower throughout the cortex, with the layer overview indicating the strongest effect in layers 2/3. In contrast, the *Ehmt1^+/−^* amygdala shows a much higher PV^+^ density already at P14. At P56, the differences are more nuanced, with a slight reduction in layers 2/3 but slightly higher densities in layers 5 and 6. In contrast to P14, the P56 PV-tdTom reporter cell density is unchanged in the amygdala. The area names are color-coded according to the Allen Brain Atlas, CCF3. In the cortex, layer 1 was blanked due to low neuron count, as were some cortical areas (minimum: 20 cells), and the layers 6a and 6b were combined into a single layer 6. Layer 4 is only plotted where it exists in the CCF3 and left blank in the other (i.e. agranular) regions. Please see supplementary table 1-4 for details.

**Figure 3:**
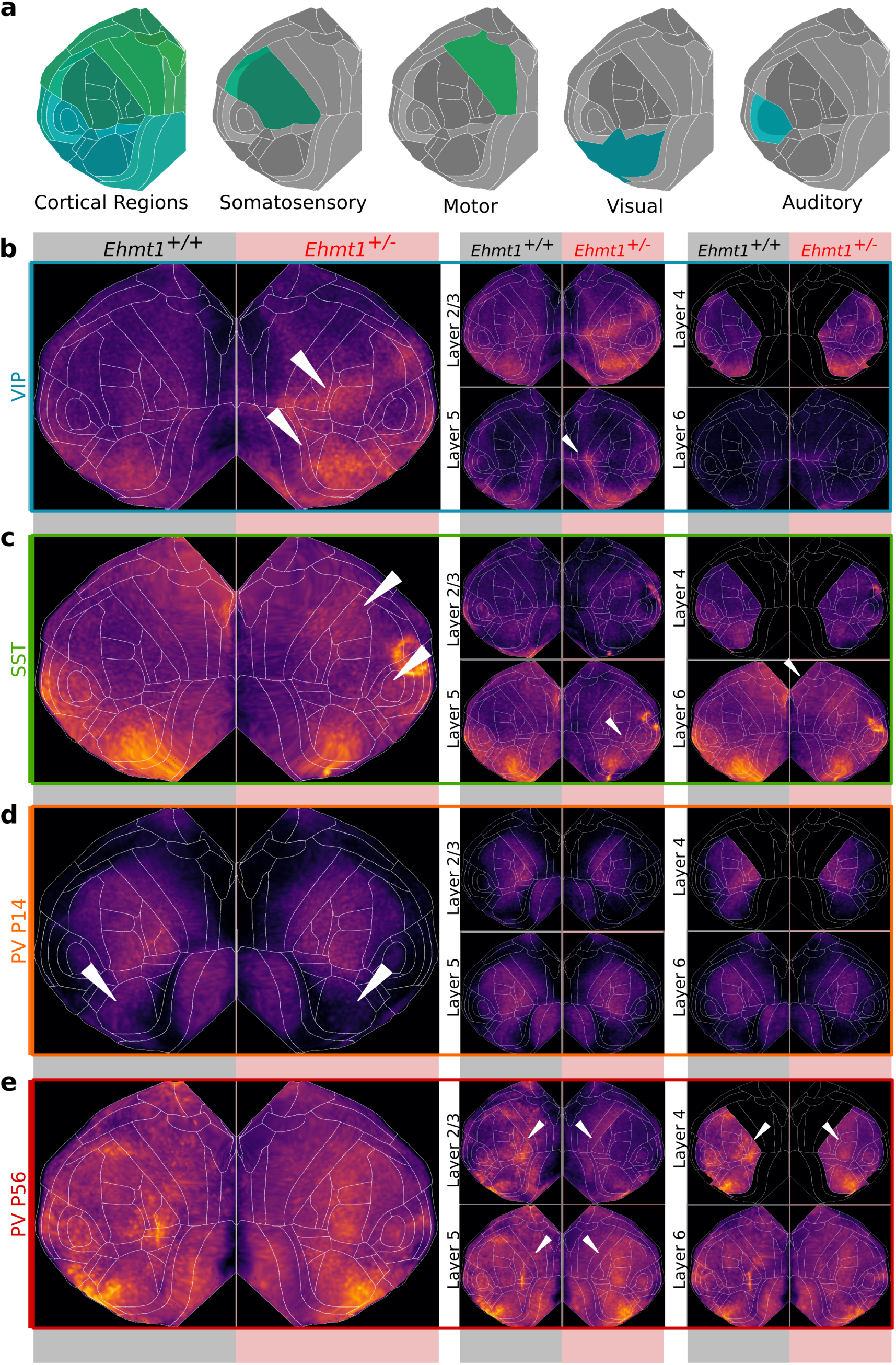
Cortical flat-map projections show cell-type differences in cortical distribution between *Ehmt1^+/+^* and *Ehmt1^+/−^*. **a)** Overview of the flat-map projection areas, colored by Allen Brain Atlas CCF3 reference color. **b)** average VIP^+^ neuron density in the cortex of *Ehmt1^+/+^* (left) and *Ehmt1^+/−^*mice (right). *Ehmt1^+/−^* cortices show two main bands of higher VIP^+^ density, one in the visual and entorhinal cortex, and another centered on the somatosensory cortex (arrowheads). Those bands appear to be centered in layers 2/3 and 4, with an additional band of higher density in layer 5 at the level of the retrosplenial cortex (arrowhead). **c)** SST^+^ cortical density. We found that SST^+^ density appears lower across the caudal end of the *Ehmt1^+/−^* cortex, roughly at the height of the auditory region (arrowheads). Similarly, the SST^+^ density appears lower at the frontal end of the *Ehmt1^+/−^*cortex, with a higher SST^+^ density in the motor cortex (arrowheads). Those changes appear to be most prominent in layers 5 and 6 (arrowheads). **d)** Adolescent PV^+^ cells are low throughout the cortex, as the cells are just starting to express PV. In accordance with the canonical sensory-driven model, we see the comparatively highest PV^+^ density in the somatosensory cortex, but also retrosplenial and orbitofrontal cortices. Interestingly, there is a small amount of PV^+^ expression in the primary visual field in *Ehmt1^+/+^*, but apparently less in *Ehmt1^+/−^* V1 (arrowheads). **e)** In adult *Ehmt1^+/−^* cortex, we find that the PV^+^ density in the somatosensory area appears shifted from layers 2/3 and 4 to layer 5 (arrowheads). Please see supplementary table 4 for details.

Next, we investigated the distribution and organization of SST^+^ neurons. SST^+^ neurons comprise ca. 30% of cortical GABAergic neurons (Bakken et al., 2021; Tremblay et al., 2016), and are thought to mostly inhibit the dendrites of pyramidal neurons (Wu et al., 2023). In our dataset, we found a reduced SST^+^ density in the *Ehmt1^+/−^* mouse brain (Fig. 2b-c), which is contrasting the overabundance of VIP^+^ neurons. Specifically, SST^+^ reduction seemed to cluster in the amygdala (basomedial amygdala BMA, intercalated nucleus of the amygdala IA, central nucleus of the amygdala CeA, anterior nucleus of the basolateral amygdala BLAa, lateral amygdala LA, cortical amygdalar area COA, piriform-amygdalar area PAA), thalamus (ventral part of the lateral geniculate nucleus LGv, ventral posterolateral nucleus VPL, ventral posterior complex VP, ventral anterior-lateral complex VAL), hippocampus (presubiculum PRE, CA1, CA2, CA3, layers 6 a/b of the ectorhinal cortex ECT6a, ECT6b) and sensory cortical areas (auditory, visual, somatosensory; Fig. 2c; Supplementary table 2). In the cortex, the reduced density was most pronounced in layers 2/3 and 4 (Fig. 2c; Fig. 3c). Finally, we examined the distribution of PV^+^ neurons across the forebrain and midbrain in *Ehmt1^+/−^* and wild-type mice at the juvenile stage (P14). PV^+^ neurons constitute the largest class of GABAergic neurons, comprising approximately 40% (Bakken et al., 2021; Gouwens et al., 2020; Tremblay et al., 2016). These neurons are the last GABAergic subtype to fully mature - marked by the expression of PV - in a process that is both brain-region and activity-dependent (Fagiolini and Hensch, 2000; Reh et al., 2020).

At the juvenile stage (P14), we observed a significant difference as *Ehmt1^+/−^* mice exhibited a higher density of PV^+^ neurons in five amygdalar subregions (posterior amygdala PA, medial amygdala MEA, dorsal endopiriform area EPd, anterior/posterior basomedial amygdala BMAa, BMAp; Fig. 2c, see Supplementary table 3 for details). This finding suggests accelerated PV^+^ maturation in the *Ehmt1^+/−^*amygdala, contrasting with our previous observations in the primary sensory cortices, where delayed PV^+^ maturation in the supragranular layers was evident at the same time (P14, Negwer et al., 2020). Whole-brain data projected onto a flattened cortical map (Fig. 3d) revealed broadly comparable PV^+^ densities between genotypes, with the highest densities observed in sensory cortices (somatosensory and auditory cortices, with minimal expression in the visual cortex). Additionally, we detected early PV^+^ expression in specific higher-order associative cortical regions, including the retrosplenial cortex (Del Rio et al., 1994) and the orbitofrontal cortex. At adulthood (P56), when PV^+^ neurons in mice are generally considered mature, we found a significant reduction in PV^+^ density in *Ehmt1^+/−^* in a secondary visual cortex (posteromedial visual area layers 2/3; Fig 2d, see Supplementary table 4 and Supplementary figure 1). Outside the cortex of *Ehmt1^+/−^* mice, we found higher PV^+^ neuron density at P56 in several subcortical areas, specifically the olivary pretectal nucleus, septofimbrial nucleus and dorsomedial nucleus of the hypothalamus (Fig. 2c, Supplementary table 4). It should be noted here that we did not quantify PV^+^ neurons in the cerebellum, given that its main cell population, i.e. Purkinje cells, express so much PV that the entire cerebellum appeared as a single, extremely bright tdTomato-positive structure.

### *Ehmt1^+/−^* amygdala shows increased PV^+^ neurons density into adulthood

The early PV^+^ hypermaturation in the amygdala (Fig. 2c) was surprising to us, because it contrasted with the slower PV^+^ maturation we had previously observed in the *Ehmt1^+/−^* sensory cortical regions (Negwer et al., 2020). Together with our finding of a reduced SST^+^ density in the amygdala at P28 (Fig. 2b), this implied a region-specific change of inhibition in the *Ehmt1^+/−^* amygdala, which prompted us to investigate this region in more detail. Using a free-floating slice staining approach with direct anti-PV immunostaining, we first investigated the density of PV^+^ neurons at the juvenile stage (P14) and in adulthood (P56). At P14, we found that PV^+^ neuron density throughout the amygdala was low, except for the posterior nucleus of the basolateral amygdala (BLAp, Fig. 4a). Still, even at P14 the neuronal density was significantly higher in the *Ehmt1^+/−^* BLAp (88/113 cells/mm², p = 0.03, t-test with 10,000 permutations, see Supplementary table 5 for details). This elevated PV^+^ neuron density in the BLAp persisted into adulthood, with similar values (Fig. 4b, 129/170 cells/mm², p = 0.045, t-test with 10,000 permutations, Supplementary table 6). The other amygdala regions showed higher PV^+^ neuron density at P56 than at P14, but no difference between genotypes (Supplementary table 5 and 6). The higher PV^+^ counts in *Ehmt1^+/−^* BLAp that persist into adulthood contrasts with our previous observations in the auditory, somatosensory, and visual cortex layers 2/3 and 4, where PV^+^ density increases slower in *Ehmt1^+/−^* than in *Ehmt1^+/+^*, eventually reaching the same level as *Ehmt1^+/+^* in all areas except for primary somatosensory cortical layer 4 (Negwer et al., 2020).

**Figure 4:**
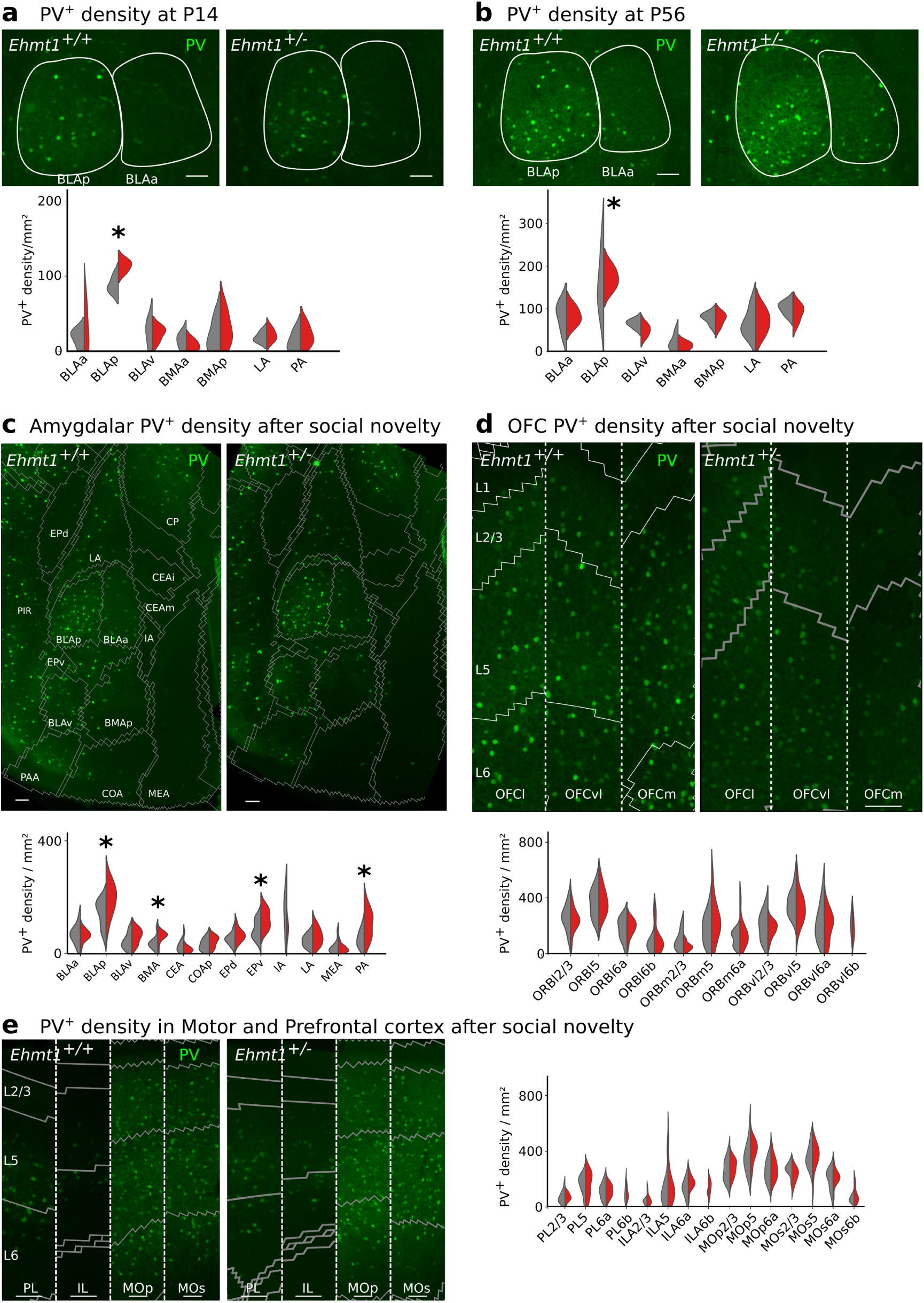
PV^+^ density in the Amygdala is higher in *Ehmt1^+/−^* mice throughout development. **a)** Quantification of PV^+^ density in different amygadalar regions at P14 in *Ehmt1^+/+^* and *Ehmt1^+/−^* mice **b)** In the Amygdala of adult naïve animals, we observed higher PV^+^ densities compared to the adolescent, still with significantly higher density in the BLAp. Representative PV^+^ immunostainings on coronal slices, with BLAp and BLAa drawn based on the Allen Brain Atlas (ca. slice 73). Scale bar: 100µm. Intensities are comparable within age groups (pixel values P14 100-2000, P56 100-5000). * = P<0.05, unpaired Student’s t-test. n=8 slices per mouse, 3-4 mice / genotype / age; Please see supplementary table 5 and 6 for details. **c)** When the Amygdala is challenged – single-housing in an enriched environment for a week, followed by social novelty exposure for 20 minutes – the PV^+^ density in the *Ehmt1^+/−^* amygdala is significantly higher in the BLAp, BMA, EPv, and PA. **d)** in the same mice, the PV^+^ density is unchanged between genotypes in the motor and frontal cortices. The relative density is highest in layer 5 of the Infralimbic, Prelimbic, primary and secondary motor cortex. **e)** Similarly, in the orbitofrontal cortex of the same mice, the PV^+^ density is unchanged between genotypes. Relative per area, the PV^+^ density is highest in layer 5. N= 48/48 slices for amygdala and 48/48 slices for frontal regions, from 12/12 *Ehmt1^+/+^ / Ehmt1^+/−^* mice, all female. Please see supplementary table 7 for details. BLA: Basolateral Amygdala, BMA: Basomedial Amygdala, LA: Lateral Amygdala, PA: Posterior Amygdala, CEA: Central Amygdala, COA: Cortical Amygdalar Area, MEA: Medial Amygdala, EP: Endopeduncular area, LHA: Lateral Hypothalamic Area, ILA: Infralimbic Area, PrL: Prelimbic Area, MOp: Primary motor cortex, MOs: Secondary Motor cortex, ORBl: Lateral Orbitofrontal cortex, ORBm: medial orbitofrontal cortex, ORBvl: ventrolateral orbitofrontal cortex.

We extended our investigation to regions closely interconnected with the basolateral amygdala (BLA), particularly those involved in social interactions (Hintiryan et al., 2021; Hintiryan and Dong, 2022). Specifically, we examined PV^+^ cell density in the amygdala, as well as in the orbitofrontal, motor, and prefrontal (prelimbic and infralimbic) cortices. We included the orbitofrontal cortex both for its reciprocal connection with the amygdala (Hintiryan et al., 2021) and because of our own finding of an elevated early PV^+^ expression already at P14 in *Ehmt1^+/−^* (Fig. 2C). For this analysis, we used brain slices from adult mice subjected to a week-long cage-enrichment and single-housing paradigm (Huang et al., 2020, 2016), which concluded with exposure to a novel social stimulus (Huang et al., 2020, 2016)(Fig. 5, supplementary Fig. 2 – 3, supplementary tables 5-7, 10). Consistent with our whole-brain screening, we observed a higher PV^+^ density in the *Ehmt1^+/−^* BLAp. Additionally, we found increased PV^+^ density in the *Ehmt1^+/−^* BMA, PA, and EPv (Fig. 4c, Supplementary table 6). These findings corroborate the earlier home-cage immunostaining results (Fig. 4a-b) and demonstrate that the accelerated PV^+^ maturation in the *Ehmt1^+/−^* amygdala, initially observed at P14 in reporter mice (Fig. 2c; supplementary Fig. 1), persists into adulthood. In contrast, PV^+^ densities in the motor, prefrontal, and orbitofrontal cortices remained unchanged between genotypes (Fig. 4d-e; Supplementary table 7 and supplementary Fig. 2). When testing for activation of those areas following a social novelty challenge by immunostaining for the immediate early gene c-Fos, we found that the *Ehmt1^+/−^* motor cortex had a higher number of c-Fos^+^ neurons after normalizing for the amount of movement (supplementary Fig. 2b; Supplementary table 7), however c-Fos^+^ density remained unchanged in both amygdala and other cortical areas (supplementary Fig. 2a-c; Supplementary table 7).

**Figure 5:**
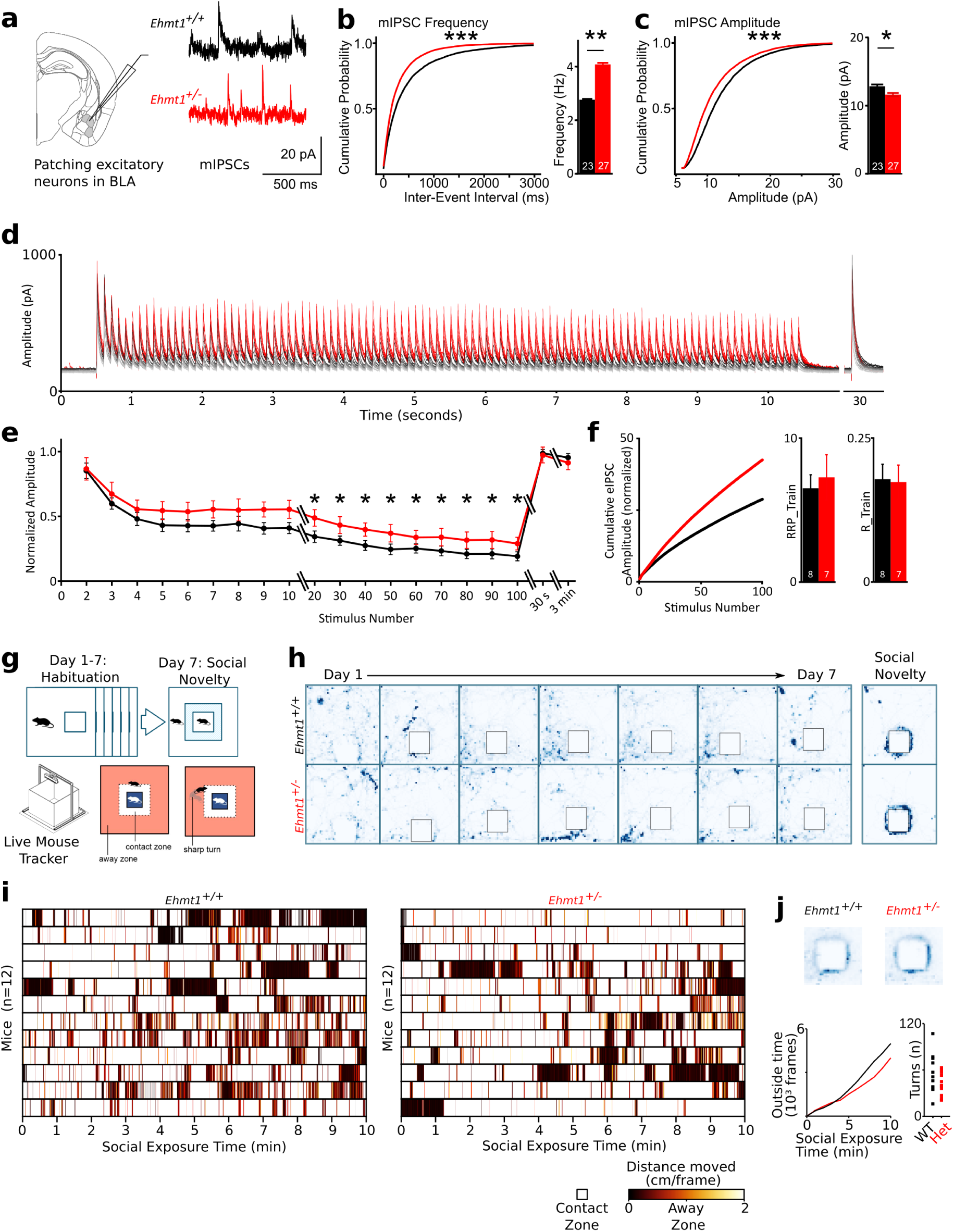
Higher GABAergic tone in the adolescent *Ehmt1^+/−^* Basolateral Amygdala, but no change in adult social behavior. **a)** Miniature Inhibitory Postsynaptic Potentials (mIPSC) obtained via whole cell voltage clamp recordings at a holding voltage of +10 mV and a pharmacological block of action potentials (TTX) and excitatory synaptic transmission (CNQX/D-AP5 for AMPA/NMDA receptors) of principal cells in the BLA of acute coronal brain sections of control (*Ehmt1^+/+^)* and *Ehmt1^+/−^* mice. **b**) Cumulative analyses for mIPSC frequency and **(c)** mIPSC amplitude. (*** = p<0.001, Kolmogorov-Smirnov test, ** = p<0.01, Student’s t test). n = 23/27 cells *Ehmt1^+/+^* / *Ehmt1^+/−^*,> 300 mIPSCs per cell. Please see supplementary table 8 for details. **d)** Representative traces for evoked GABAergic postsynaptic currents (eIPSCs) measured at a holding potential of 0 mV with a pharmacological block of NMDA receptors. Shown here are traces from 10 sweeps from a representative cell for each *Ehmt1^+/+^* (black) and *Ehmt1^+/−^* (red). **e)** Relative eIPSC amplitude normalized to the first peak of the sweep. (* = p<0.05, unpaired Student’s t-test) n = 8/7 cells *Ehmt1^+/^*^+^ / *Ehmt1^+/−^*, each 10 sweeps with 100 stimuli each. Please see supplementary table 9 for details. **f)** Cumulative (normalized) amplitudes over the train and estimated Readily Releasable Pool (RRP) and Release Probability (p>0.05, unpaired Student’s t-test). n=8/7 cells *Ehmt1^+/+^* / *Ehmt1^+/−^* 10 sweeps with 100 stimuli each. **g)** Experimental paradigm: An “experimental mouse” of either genotype was single-housed for 7 days in a LiveMouseTracker cage (50×50×35 cm), we measured their movement in 2 consecutive sessions of 10 minutes each day. **h)** On the last session of the last day, a “stranger mouse” was placed in the mini-cage at the center of the LMT cage. We then quantified the time and movement of the experimental mouse in the “outside zone”, when the head was more than 5 cm removed from the mini-cage. We found no difference in movement (**i**), time in the outside zone or the number of turns (**j**) between genotypes. n=12/12 *Ehmt1^+/+^ / Ehmt1^+/−^* mice, all female adults. Please see supplementary table 10 for details.

### Principal neurons in the *Ehmt1^+/−^* amygdala receive elevated GABAergic input

We next investigated functionally whether the higher density of PV^+^ neurons in the adolescent *Ehmt1^+/−^* amygdala translates to an increased inhibitory drive. To this end, we measured GABAergic miniature inhibitory postsynaptic currents (mIPSCs) in BLA principal neurons in acute *ex vivo* brain slice preparations at P12-14 (Fig. 5a). mIPSCs represent spontaneous single-vesicle releases of GABA in the absence of network activity. Their frequency is considered to reflect the number of perisomatic GABAergic synapses, while their amplitude correlates with average synaptic strength (Dervinis and Major, 2022; Liao et al., 1995; Segal, 2010). In *Ehmt1^+/−^* mice, we observed a combination of higher mIPSC frequency (Fig. 5b; IEI 517 ms/ 312 ms; p = 0.007, nested ANOVA, 23/27 cells with >300 mIPSCs each) and lower amplitude (Fig. 5c; 12.8 pA / 11.6 pA; p= 0.0420, nested ANOVA, 23/27 cells with > 300 mIPSCs each; Supplementary table 8), indicating an increased number of GABAergic synapses with reduced synaptic strength.

To determine whether the increase in spontaneous GABA release is associated with enhanced stimulated GABAergic signal transmission, we employed electrical stimulation consisting of a 10 Hz pulse train over 10 seconds, designed to deplete presynaptic releasable vesicle pools and test steady-state transmission (Brager et al. 2003). The initial GABAergic evoked inhibitory postsynaptic current (eIPSC) amplitudes were comparable across genotypes (Fig. 5d; 398±22 pA / 363±24 pA, *Ehmt1^+/+^/Ehmt1^+/−^*, p = 0.98). Over the course of the stimulus train, the relative eIPSC amplitudes (normalized to the first response of each sweep) dropped by approximately 50% during the first 10 stimuli in both genotypes (Figs. 5d–e). However, *Ehmt1^+/−^* neurons displayed significantly higher sustained eIPSC amplitudes throughout the remaining stimuli (Fig. 5e, 21% / 31% by pulse 90, n = 8/7 *Ehmt1^+/+^/Ehmt1^+/−^* cells, Supplementary table 9). Following a 30-second recovery interval, eIPSC amplitudes recovered to the level of the initial peak in both genotypes (Fig. 5e), suggesting that the observed differences were not due to a larger readily releasable pool (RRP) of synaptic vesicles per synapse (Neher, 2015), but rather a higher number of synapses contributing to steady-state transmission.

Using linear extrapolation of the cumulative amplitude plots (Elmqvist and Quastel, 1965; Neher, 2015) further confirmed that the RRP and release probability were unchanged between genotypes (Fig. 5f; RRP 6.5±0.9 / 7.2±1.5 p = 0.67; Release Probability 0.17±0.03 / 0.17±0.03, p = 0.89; n=8/7 *Ehmt1^+/+^ / Ehmt1^+/−^* cells, see Supplementary table 9). Collectively, our results point to a larger number of GABAergic synapses in the BLA of *Ehmt1^+/−^* mice. While individual synaptic strength was lower, the increased synapse count facilitated higher sustained GABAergic transmission. This aligns with our finding of increased PV^+^ neuron density in the BLA.

### Subtle changes in *Ehmt1^+/−^* behavior

The change in amygdalar inhibitory network composition – increased PV^+^ density, higher mIPSC frequency and higher sustained GABAergic transmission, but lower mIPSC amplitude – led us to hypothesize that the amygdalar circuit might be at a homeostatic activity setpoint, where the *Ehmt1^+/−^* circuit may receive more input, but local activity is dampened by stronger PV^+^ connectivity. We therefore expected this circuit to be less stress-tolerant and therefore expected that a strong social stimulus following social isolation would show changes in behavioral activity and neuronal activity, as described in the social novelty paradigm from Huang and colleagues (Huang et al., 2020, 2016): week-long single housing followed by novel social exposure to a “stranger mouse” (Fig. 5g) generated neuronal high activity levels - measured with c-Fos^+^ immunostainings as a proxy - in both BLA and OFC. Based on previous three-chamber experiments in *Ehmt1^+/−^* (Balemans et al., 2010), we expected the *Ehmt1^+/−^* mice to have a “social persistence phenotype” where the *Ehmt1^+/−^* mice would spend more time with the stranger mouse than elsewhere. We adapted this paradigm to the LiveMouseTracker platform (De Chaumont et al., 2019), with week-long single housing in a 50×50 cm home cage with a small, locked 10×10×10 cm mini-cage with perforated-metal sides and plexiglass top placed inside. During the first 7 days, we habituated the mice to the LiveMouseTracker recordings by placing the cage under the recording setup (during the dark period) and recording two sessions of 10 minutes each. On the last session of the last day, we placed an unrelated, wild-type “stranger mouse” into the mini cage, and recorded 10 minutes of the experimental mouse’s movements around the cage. We designated a 5 cm zone around the cage as “contact zone”, and the rest as “outside zone”. When analyzing the recordings, we found large individual differences in the reactions, but no clear difference between genotypes in movement speed (Fig. 5i), time in the outside zone and number of turns (Fig. 5j; N = 12/12 female *Ehmt1^+/+^* / *Ehmt1^+/−^* mice; Supplementary table 10). When we investigated the brains of those mice for c-Fos^+^ activity, we did not find any difference in amygdala (supplementary Fig. 2a) or orbitofrontal regions (supplementary Fig. 2c), with a single exception in the motor cortex (supplementary Fig. 2b; Supplementary table 7). Similarly, we did not find differences in behavior of conditional *Ehmt1* heterozygous knockouts in PV^+^ neurons (supplementary Fig. 3a; Supplementary table 10).

However, using the LiveMouseTracker to observe group behavior in familiar groups (cage mates since at least weaning) over a 22-hour period (supplementary Fig. 3b), we observed a change in the diurnal activity pattern in *Ehmt1^+/−^* mice, with a lower activity peak after the onset of the dark period (supplementary Fig. 3c). Further investigation revealed that this difference was entirely driven by males, which showed a reduction of ca. 10% in their movement in the first two hours of the dark period (the active time for nocturnal animals such as the mouse; supplementary Fig. 3d-e. n = 14/14 *Ehmt1^+/+^/Ehmt1^+/−^* females, and 14/14 *Ehmt1^+/+^/Ehmt1^+/−^* males).

To conclude, while there are prominent cellular and synaptic phenotypes in the *Ehmt1^+/−^* brain, the behavioral phenotypes we measured – in our paradigms – appear to be comparatively subtle.

## Discussion

In this study, we investigated the effects of *Ehmt1* haploinsufficiency on GABAergic neuron subclasses across multiple brain regions at different developmental stages. Our findings reveal distinct changes in the density of SST^+^, PV^+^, and VIP^+^ neurons, highlighting regional specificity in the impact of *Ehmt1^+/−^* on GABAergic circuits. These alterations may have broad implications for GABAergic inhibitory network function, particularly in the context of fear processing and sensory integration.

### Does enhanced amygdalar inhibitory drive explain the *Ehmt1^+/−^* fear phenotype?

The most pronounced changes were observed in the cortex and amygdala. In the *Ehmt1^+/−^* cortex, VIP^+^ neuron density was markedly elevated. In contrast, in the BLA we observed a significant reduction in SST^+^ neuron density, accompanied by an increase in PV^+^ neuron density, evident already at the juvenile stage. The increased PV^+^ density in the amygdala was associated with heightened spontaneous and evoked GABAergic transmission. Our whole-cell patch-clamp approach overrepresents input close to the pipette on the soma, so we might have underestimated a potential reduction of inhibitory drive onto the dendrites, which is specifically generated by SST^+^ neurons. The fact that we saw no change in amygdalar c-Fos^+^ expression following a social novelty challenge could mean that the circuit activity is maintained by a homeostatic mechanism. However, a reduction of SST^+^-mediated dendritic inhibition with a strengthening of PV^+^-mediated perisomatic inhibition could reduce dendritic computation capability and reduce input selectivity. Indeed we may be seeing a first indication of this in the mIPSC kinetics (supplementary Fig. 7), where *Ehmt1^+/−^* has a slower rise but faster decay time, which potentially indicates more fast GABAergic synapses (presumably perisomatic PV^+^-mediated) and fewer slow GABAergic synapses (presumably distal SST^+^-mediated). This is in line with a recent computational model proposed by Cattani and colleagues (Cattani et al., 2024). Similarly, our cortical findings in e.g. somatosensory cortex of higher layer 5-6 VIP^+^ density and lower layer 6 SST^+^ density would converge on a reduced SST^+^ dendritic inhibition, both due to lower SST^+^ density as well as higher VIP^+^-mediated inhibition of the remaining SST^+^ neurons (Preuss et al., 2026).

Within the BLA the shift in GABAergic inhibition from dendritic to more somatic synapsing domains could in turn influence local oscillatory activity and oscillatory coupling to other regions. Recent work from Pankaj Sah’s group has shown that PV^+^ neurons in the BLA can reliably evoke disynaptic feedback excitation onto themselves (Spampanato et al., 2016). Activity in a single BLA PV^+^ neuron is sufficient to create sharp-wave ripple activity in *ex vivo* brain slice preparations (Perumal et al., 2021), which translates to gamma oscillations *in vivo* (Feng et al., 2019). Elevated gamma oscillations in the BLA could strengthen fear memory encoding and stability, as gamma-band synchrony has been implicated in fear memory entrainment and extinction learning (Courtin et al., 2014; Trouche et al., 2013). Furthermore, fear behavior is also modulated by SST^+^ neurons in the central amygdala (Sun et al., 2023), and SST^+^ neurons control beta-synchrony between amygdala and hippocampus, a connection in which small timing shifts modulate fear behavior (Jackson et al., 2024). This is especially interesting in the light of the fear behavior phenotype of the *Ehmt1^+/−^* mice, which show higher anxiety in an elevated plus maze (Balemans et al., 2010) and a complete lack of fear extinction and impaired glutamatergic synaptic release at the CA3-CA1 connection (Balemans et al., 2013). The lack of extinction could be mediated by a dysregulated immune response in *Ehmt1^+/−^* mice (Yamada et al., 2021), as hippocampal microglia have recently been shown to be involved in extinction learning, where they are recruited to temporarily silence fear engram neurons in the dentate gyrus (Liu et al., 2026). That said, the expression of the anxiety behaviour might still be context-dependent, as our current study - with its stress-reducing home-cage imaging paradigm - only found modest behavioural differences. In KS patients, anxiety is also common in ca. 45% of patients (Vermeulen et al., 2017a). Our findings suggest that the increased PV^+^ density (with concurrent reduced SST^+^ density) and enhanced inhibitory drive in the BLA could contribute to more stable gamma-band oscillatory activity, which would align with the behavioral phenotypes observed in *Ehmt1^+/−^* mice. Both GABAergic neuron populations as well as the BLA itself can be subdivided further, both by projection pattern (Hintiryan and Dong, 2022) as well as by CCK-defined functional subnuclei with distinct excitatory populations (Reéb et al., 2025).

### Are sleep disruptions in KLEFS1 linked to subcortical PV^+^ and VIP^+^ populations?

In KLEFS1 patients, sleep disruption is a common occurrence and is in fact predictive of severe regression (Vermeulen et al., 2017b). This matches with both the behavior and cellular changes we observed in *Ehmt1^+/−^* mice. Specifically, the only behavioral phenotype we observed in our mice was a change of circadian activity in males (supplementary Fig. 3e), which might be indicative of a disrupted sleep homeostasis. Furthermore, in our whole-brain VIP^+^ screening, we saw significantly lower VIP^+^ neuron densities in the midbrain reticular nucleus. VIP^+^ signaling in the midbrain reticular nucleus has been directly linked to sleep, specifically the onset of REM sleep (Kohlmeier and Reiner, 1999). We also observed significantly higher PV^+^ densities in two subcortical regions are involved in sleep homeostasis: the olivary pretectal nucleus is thought to encode circadian light-darkness levels (Prichard et al., 2002) in GABA-dependent slow oscillations (Szkudlarek et al., 2012, 2008). Specific GABAergic populations also code for activity levels during the mouse’s active phase in the dorsomedial hypothalamus (Liu et al., 2023) and in the dorsal Raphe nucleus under stress (Ren et al., 2024). The Suprachiasmatic nucleus has also been directly implicated in light-dependent diurnal activity regulation, specifically via VIP^+^ neurons (Kahan et al., 2023; Todd et al., 2020; Wang et al., 2026). Unfortunately in our samples, the SCN is not present in all samples as it is very close to the midline (supplementary Fig. 4 a). A dysregulation of the specific oscillatory patterns in those regions might be linked to the change in diurnal activity patterns we observed in *Ehmt1^+/−^* male mice (supplementary Fig. 3e) and the disrupted sleep observed in KS patients (Vermeulen et al., 2017b). To gain a more robust understanding, future studies should use whole brains (not just hemispheres) to better preserve the details along the midline, and use larger groups so that male and female mice can be compared as separate groups.

### Cortical PV^+^ neurons: Subtle changes obscured by tdTomato reporter?

Our iDISCO+ data did not reveal changes in the PV^+^ neuron density in the cortex, which contrasts with our previous study (Negwer et al., 2020) that used detailed slice stainings and found slightly delayed PV^+^ maturation in three main cortical sensory areas (auditory, somatosensory and visual) layers 2/3 and 4. This may be due to the large number of cortical areas we are comparing here, as there are no statistically significant differences in the PV^+^ iDISCO+ data after multiple-testing correction (Supplementary table 3, Figure 2c). However, this could also be explained by methodological differences: Whereas the previous study used direct anti-PV immunolabeling, our current study used a genetic labeling strategy with a Cre-dependent tdTomato reporter. The reporter line provides a binary staining, where an activation of the *Pvalb* gene at any point during development will drive a strong and permanent expression of tdTomato via the CAG promoter and render the cell permanently fluorescent, independent of its current PV protein expression. In contrast, our anti-PV staining on slices (Negwer et al., 2020 and Fig. 4) are capturing the momentary PV expression that is expected to rise gradually and vary depending on activity. In consequence, the reporter line might lead to an overestimate of PV^+^ cells both by cell lineage (some cells activating the *Pvalb* promoter early during development but not expressing PV later, for example some SST^+^ subpopulations in Gouwens et al., 2020), and by timing mismatch (tdTom being expressed faster and stronger than the more gradual and activity-dependent rise in Parvalbumin expression).

We see evidence of the latter in the cortex around P14, where the *Pvalb* promoter has already been active (and therefore tdTom is being expressed) but PV protein expression is only gradually rising. In contrast, in the P14 Amygdala with its higher expression levels, we see an agreement between reporter expression (Fig. 2) and Parvalbumin staining levels (Fig. 4), in that both counts are higher in *Ehmt1^+/−^* brains.

Future studies should consider using either a stable marker for (future) PV^+^ cells that is expressed before PV (e.g. PGC1a; Moissidis et al., 2025), stain directly for PV or other markers of PV maturation such as perineuronal nets, to match the temporal resolution of their staining to the temporal requirements of their research question.

### Possible developmental origins of GABAergic neuron population shifts: Progenitor pool dynamics or cell survival?

In the cortex, our findings of a lower SST^+^ and higher VIP^+^ neuron density could be explained by a shift in the progenitor population. The precursors to GABAergic neurons are generated in temporary structures in the developing mouse brain called ganglionic eminences, specifically the medial ganglionic eminence (MGE) for SST^+^ neurons and caudal ganglionic eminence (CGE) for VIP^+^ neurons (Kessaris et al., 2014; Kessaris and Denaxa, 2023). The specification of the MGE/CGE boundary is governed by gradients of transcription factors (Kessaris et al., 2014, their table 1) and epigenetic modifiers (Allaway et al., 2021; Mossink et al., 2021). *Ehmt1* haploinsufficiency could therefore lead to a shift in the MGE/CGE boundary, leading to a larger progenitor pool for VIP^+^ neurons at the expense of SST^+^ progenitors. Furthermore, each of these cell classes can be subdivided further into subclasses, by laminar localization, gene expression, or morphotype (Gouwens et al., 2020; Fisher et al., 2025). For example, in the layers 5-6 of the *Ehmt1^+/−^* somatosensory cortex, we find a significantly higher VIP^+^ and lower SST^+^ density (Fig. 2C). Especially layer 6 SST^+^ cells are enriched in the long-range projecting subtype of SST^+^ cells (Fisher et al., 2025), which might be especially impacted in the *Ehmt1^+/−^* mouse. In contrast, the SST^+^ densitites in the upper layers seem mostly unaffected. This is where the SST^+^/CR^+^ subtype is found, which is thought to derive from the Nkx6.2^+^ MGE/CGE boundary (Fogarty et al., 2007). Future studies could use both layer 6 SST^+^ LRP and layer 2/3 SST^+^/CR^+^ cell classes as especially promising targets for changes in the *Ehmt1^+/−^* cortical inhibitory lineup.

In the amygdala however, we found premature PV^+^ maturation and higher PV^+^ density even into adulthood. This is unlikely to be due to a change in progenitor populations, as the amygdalar PV^+^ progenitors seem to be generated from the same MGE-derived progenitor pool as cortical PV^+^ neurons (Aerts and Seuntjens, 2021). The amygdalar differences could be explained by activity-dependent postnatal cell survival (Wong et al., 2018) for PV^+^ and SST^+^ cells, if this concept can be extended from the cortex to the amygdala. This would fit with both the higher PV^+^ density we observe in the adult amygdala and the premature PV^+^ expression at P14, as PV^+^ maturation is activity-dependent (Moissidis et al., 2025).

## Conclusion

To conclude, we conducted the first whole-brain GABAergic neuron survey in an ASD mouse model. We found cell-type and region-specific effects of *Ehmt1* haploinsufficiency (Figure 6), with key effects in the cortex (VIP^+^ higher, SST^+^ reduced in *Ehmt1^+/−^*), amygdala (PV^+^ higher, SST^+^ reduced) and sleep-related subcortical regions (VIP^+^ and PV^+^ higher). We investigated the amygdala in greater detail and confirmed a higher PV^+^ density already at P14 that lasts into adulthood. This elevation in PV^+^ density was reflected in a higher spontaneous and evoked GABAergic transmission. However, we found no change in post-behavioral amygdalar or orbitofrontal neuronal activation (c-Fos^+^ density), which was matched by the unchanged behavior, possibly indicating that the higher PV^+^ density is a circuit-level homeostatic mechanism to keep the amygdalar activity at an innate set point. We did however find a reduction in diurnal activity in *Ehmt1^+/−^* male mice, which could relate to the changes in VIP^+^ and PV^+^ density in sleep-related regions we identified with our whole-brain screening and point towards a circuit mechanism for the sleep problems identified in KLEFS1 patients. To our knowledge, our study is the first whole-brain GABAergic screening in an ASD mouse model, and the surprising finding of an early PV^+^ maturation in the *Ehmt1^+/−^* amygdala shows that a broad initial screen can light the way to interesting regions that can then be investigated in detail.

**Figure 6:**
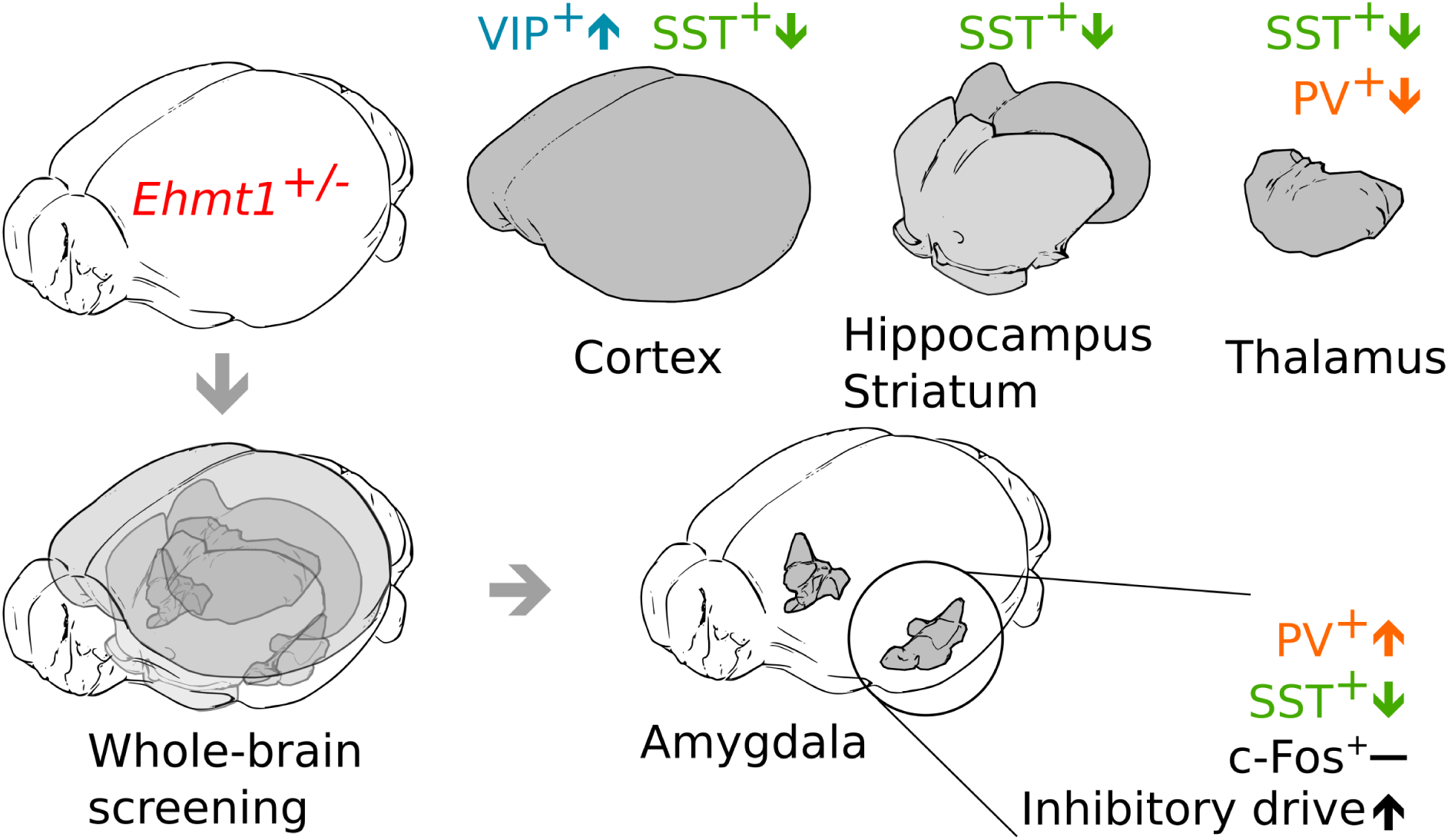
Summary figure. Using an unbiased, whole-brain clearing protocol we found that the *Ehmt1^+/−^* cortex contains more VIP^+^ cells, but fewer SST^+^. We also found that the hippocampus and striatum contain fewer SST^+^ cells. The *Ehmt1^+/−^* thalamus contains fewer SST^+^ and (early) PV^+^ cells. The amygdala, surprisingly, contains fewer SST^+^ cells but more PV^+^ cells. We confirmed the latter with slice immunostainings and electrophysiological measurements. We found that the juvenile *Ehmt1^+/−^* BLA has a higher GABAergic transmission, but no change in c-Fos^+^ upon a social novelty challenge.

## Material and Methods

### Transgenic mice

The mice crossing was performed as described in (Negwer et al., 2022). We used reporter mice expressing a subpopulation-specific Cre Recombinase driver (see below) and a floxed tdTomato reporter (ai14, see below), on a mixed C57/Bl6J background. We crossed *Cre* homozygotes with *tdTomato^flox/flox^ / Ehmt1^+/−^*, and used the first-generation offspring that was heterozygous for both Cre and tdTomato-flox reporter expression. Specifically, we used the following mouse lines:

- *PV-Cre*: B6.129P2-Pvalb^tm1(cre)Arbr^/J, RRID:IMSR_JAX:017320
- *SST-Cre*: B6J.Cg-Sst^tm2.1(cre)Zjh^/MwarJ, RRID:IMSR_JAX:028864
- *VIP-Cre*: B6J.Cg-Vip^tm1(cre)Zjh^/AreckJ, RRID:IMSR_JAX:031628
- *tdTom-flox*:B6.Cg-Gt(ROSA)26Sor^tm14(CAG-tdTomato)Hze^/J,RRID:IMSR_JAX:007914
- *Ehmt1^+/−^*: Ehmt1^tm1Yshk^ RRID:MGI:3575661
- *Ehmt1^flox^*: C57BL/6Smoc-Ehmt1^tm1(flox)Smoc^ RRID:IMSR_NM-CKO-190020

Mice were kept as described previously (Negwer et al., 2020). In brief, we used mice from both sexes for our study. Animals were kept in wire-top cages (type III) and group-housed with the with the nest until weaning (P21-25), or and afterwards group-housed (3-6 animals, littermates, both genotypes, segregated by sex). Animals had access to rodent chow and water *ad libitum* and were kept on a 12h/12h light-/dark cycle (lights on at 07:00 AM). Animal experiments were conducted in conformity with the Animal Care Committee of the Radboud University Nijmegen Medical Centre and the National Committee on Animal Experiments (CCD), The Netherlands, and conform to the guidelines of the Dutch Council for Animal Care and the European Communities Council Directive 2010/63/EU (Project RU-DEC 2017-0048, permit AVD10300 2019 7746).

### Brain clearing

We performed iDISCO+ stainings as described in (Negwer et al., 2022, 2020). In short, we processed one hemisphere per brain following the iDISCO+ histochemistry protocol (Renier et al., 2016) as described for adult brains, with all buffers according to the protocol and all incubation steps taking place on a shaker/rotor, in 5 ml screw-top Eppendorf tubes, and lasting 1h unless mentioned otherwise. For details, please see the protocol at protocols.io (https://dx.doi.org/10.17504/protocols.io.36wgq77m5vk5/v1). Whole mouse brain hemispheres were dehydrated in a methanol gradient (from 20% to 100%), bleached in 5% H_2_O_2_ in methanol at 4°C overnight, then rehydrated. Hemispheres were subsequently permeabilized for 5-7 days at RT, blocked for 5-7 days at 37°C, then incubated with primary antibodies (Rabbit Anti-RFP, Rockland 600-401-379, 1:2000, 2 ml / sample) for 6 days at 37°C. Subsequently, brains were washed 5×1h + 1x overnight at RT, and incubated for 7 days with secondaries (Goat anti-rabbit Alexa-568, Invitrogen A11036, or Goat anti-Rabbit Alexa-647, Invitrogen A-21245 1:500, 2 ml / sample) at 37°C. Following 5x 1h + 1x overnight washing at RT, samples were dehydrated in a methanol gradient, then twice more in 100% methanol, 66% DCM/33% methanol, 2x 15 min 100% DCM, and finally cleared in 100% Dibenzyl Ether (DBE, Sigma) in airtight glass vials. Brains were typically transparent within 2h, and completely cleared overnight. Alexa fluorescence in the samples remained at useable levels for at least 12 months of storage in DBE and over (maximally) 3 imaging rounds.

| Name | Dilution | Supplier | Cat# | RRID | Used for |
| --- | --- | --- | --- | --- | --- |
| Goat anti-Rabbit Alexa 647 | 1:500 | Invitrogen | A-21245 | RRID:AB_2535813 | VIP-tdTom |
| Goat anti-Rabbit Alexa 568 | 1:500 | Invitrogen | A-11036 | RRID:AB_10563566 | PV-tdTom (P14 / P56), SST-tdTom |
| Rabbit anti-RFP | 1:2000 | Rockland | 600-401-379 | RRID:AB_2209751 | All iDISCO+ |
| Mouse anti-Parvalbumin | 1:1000 | Swant | 235 | RRID:AB_10000343 | Slice stainings |
| Rabbit anti-cFos | 1:500 | Synaptic Systems | 226 008 | RRID:AB_2891278 | Slice Stainings |

### Light-sheet microscopy imaging

The cleared samples were imaged on a Miltenyi LaVision Ultramicroscope II Light-sheet microscope outfitted with a NTK Photonics white-light laser and filter sets for 488 nm, 568 nm, and 647nm, imaged through a long-working distance objective (Miltenyi) at 1.1x magnification (effective 2.2x, NA 0.1), and recorded with an Andor Neo 5.5 cooled sCMOS camera. We imaged with a 480 nm signal for autofluorescence for alignment with the Allen Brain Atlas, and 560 nm (or 640nm) to record the TdTomato-Alexa signal. We used a single light-sheet from one side at 0.54 NA, scanning at 2.95 / 2.95 / 3 µm x/y/z resolution (3 µm z-steps) with the “horizontal focus” method and 17-18 horizontal focus steps. The sample was imaged submerged in DBE in sagittal configuration, and the entire cortex fit inside a single field of view (x/y), with a typical brain producing ∼1600 z-planes of 3 µm each.

### Light-sheet stack processing with FriendlyCearMap

We used FriendlyClearmap (Negwer et al., 2022) to analyze the resulting image stacks. In brief, this pipeline is an adapted version of ClearMap (Renier et al., 2016) that we rewrote in Python 3.7, added machine-learning based cell segmentation with Ilastik (Berg et al., 2019), and a landmark-based atlas registration with BigWarp (Bogovic et al., 2016) and Elastix (Klein et al., 2010). We generated a custom training set in Ilastik for each set of brains (VIP, SST, PV P14 and PV P56), which we then used to segment all brains from that set. We then manually generated ca. 50-100 landmarks on the autofluorescence channel and the CCF3 reference atlas for each brain using BigWarp, and used Elastix (via FriendlyClearmap) to transform the brain’s image stacks to the reference atlas. We used the age-adapted CCF3 atlas variants from (Newmaster et al., 2020) for SST^+^ and VIP^+^ brains (P28) and juvenile PV^+^ (P14) and the Allen Brain Institute’s CCF3 atlas (Oh et al., 2014; Wang et al., 2020), downloaded from the Scalable Brain Atlas page (Bakker et al., 2015) for the P56 PV^+^ mice. FriendlyClearmap generated a detailed per-region cell count, for which we calculated significant differences with a nonparametric Mann-Whitney *U* test not corrected for multiple comparisons. We ran all cell segmentation, landmark generation, and postprocessing on a single laptop (AMD Ryzen 4600H, 64GB RAM, 1TB SSD) running Linux (Ubuntu 24.04 LTS) and used FriendlyClearMap inside a docker container. For a detailed description, please refer to protocols.io for FriendlyClearMap (https://dx.doi.org/10.17504/protocols.io.yxmvmn9pbg3p/v3, https://dx.doi.org/10.17504/protocols.io.eq2lynnkrvx9/v2).

### Landmark selection for atlas registration

We selected the landmarks in BigWarp, as explained in the Friendlyclearmap documentation (https://dx.doi.org/10.17504/protocols.io.eq2lynnkrvx9/v2), section 10; see also supplementary figure 5 of this paper). We selected 50-100 landmarks between our downsampled brain stack and the respective atlas stack. Specifically, we paid special attention to the following regions for landmark selection: Hippocampus (CA1, DG), Cortex (pia and white matter in an imagined column in e.g. S1 barrel field and the frontal pole of the PFC, visual cortex, motor cortex and retrosplenial cortex), thalamic reticular nucleus, fiber bundles, hypothalamus, amygdala. We validated the success of the atlas registration manually by Elastix’s tiff output feature, overlaying the warped brain stacks onto the atlas in Fiji. If the overlap was deemed insufficient, we made a new set of landmarks, paying more attention to the previously misaligned regions.

### Single-cell electrophysiology

We performed single-cell electrophysiology as described in (Frega et al., 2019; Negwer et al., 2020). We used litter-matched *Ehmt1^+/+^* and *Ehmt1^+/−^* mice of adolescent age (postnatal day P14-16). In brief, animals were deeply anesthetized with isoflurane, then quickly decapitated. 350-µm-thick coronal slices were cut using a microtome (HM650V, Thermo Scientific) in ice-cold “cutting and storage” ACSF containing 87mM NaCl, 11mM Glucose, 75 mM Sucrose, 2.5 mM KCl, 1.25 mM NaH2PO4, 0.5 mM CaCl2, 7mM MgCl2, and 26mM NaHCO3, continuously oxygenated with 95% O2/5% CO2. Slices were incubated for 1 h at 32 °C, after which they were allowed to cool down to room temperature. The slices were then transferred to an upright microscope (Olympus) fitted with 2.5x and 40x water-immersion objectives and enhanced infrared illumination (DGC, Luigs & Neumann), and incubated in “recording” ACSF (124 mM NaCl, 10 mM Glucose, 3 mM KCl, 1.25 mM NaH2PO4, 2mM CaCl2, 1mM MgCl2, and 26 mM NaHCO3) at 30 °C.

For measuring miniature IPSCs (mIPSCs), slices were incubated in recording ACSF with 1 µM TTX (Tocris 1069) to block action potentials, and 100 µM D-AP5 (Tocris 0106) and 5.2 µM CNQX (Tocris 1045) to block excitatory transmission. Slices were allowed to adjust to the conditions for at least 10 minutes prior to patching. Pyramidal cells in BLA were identified based on their perisomatic appearance at 40x magnification, and whole-cell patch-clamp recorded using 3–6MΩ borosilicate pipettes filled with a Cesium-based intracellular solution containing 115 mM CsMeSO3, 20 mM CsCl2, 10 mM HEPES, 2.5 mM MgCl2, 4mM Na2ATP, 0.4 mM NaGTP, 10mM Na-Phosphocreatine, 0.6 mM EGTA, and 5 mM QX-314 to block action potentials. Cells were recorded at a minimum depth of 30 µm from the slice surface to minimize cutting artefacts. The cells were recorded in voltage-clamp, controlled by an SEC 05-LX amplifier (NPI electronics, Tamm, Germany), low-pass filtered at 3 kHz and sampled at 20 kHz with a Power-1401 acquisition board and Signal software (CED, Cambridge, UK). The holding voltage (V_h_) was adjusted to +10 mV, and miniature inhibitory postsynaptic currents (mIPSCs) were recorded over 10 min per cell. Subsequently, the traces were exported from Signal as text files, saved as .abf with Clampfit and then loaded into MiniAnalysis. mIPSCs were identified manually, with an amplitude threshold of 8 pA, at least 300 events per cell.

To determine evoked GABA release, slices were incubated in recording ACSF with 100 µM D-AP5 (Tocris 0106) and 5.2 µM CNQX (Tocris 1045) added to block AMPA and NMDA-mediated currents. A tungsten bipolar stimulation electrode (CE2C55, FHC) coupled to an SD9 stimulator (Grass Instruments, RI, USA) or an ISO-STIM 01M (NPI electronics, Tamm, Germany) was inserted into the BLA, and principal neurons were patched next to the bipolar electrode (<200 µm lateral distance) as described above.

GABAergic currents were measured at V_h_ = 0 mV. Stimulus duration (1-2 ms) and voltage were adjusted for a postsynaptic GABAergic amplitude of ∼400 pA. We measured GABAergic release probabilities via a 10 Hz rundown protocol. In this protocol, we recorded the cell at V_h_ = 0 mV and evoked a train of 100 stimuli at 10 Hz to deplete the GABAergic vesicle pool. We tested for recovery with an additional stimulus after 30s. Inter-sweep interval was 180 s. We exported the data as text files and determined peak amplitude with a custom Python script. We normalized the peak amplitude to the first peak in the sweep’s stimulus train. We calculated the readily releasable pool and the release probability by curve-fitting on a plot of the individual response amplitude against cumulative amplitude (Elmqvist and Quastel, 1965; Neher, 2015). To get the RRP, normalized amplitudes (y-axis) were plotted against normalized cumulative amplitudes (x-axis). The steep initial change was extrapolated linearly, and the x-axis intercept gives the estimate of the readily releasable pool (RRP). The slope gives the estimate of the compound release probability (PR) (n = 8/7 *Ehmt1^+/+^/Ehmt1^+/−^* cells).

### Brain slice immunohistochemistry

Brain sections were stained similar to the protocol described in (Negwer et al., 2020). In short, mice were deeply anesthetized with isoflurane and either decapitated (for Fig. 4a-b) or transcardially perfused with ice-cold PBS followed by 4% PFA in PBS (Fig 4c-e, supplementary Fig. 2). In both cases, the brains were then extracted out of the skull and postfixed overnight in 4% PFA in PBS at 4°C, then rinsed in PBS and stored in PBS + 0.005% Sodium Azide until processing. The brains were then embedded in agarose blocks and cut coronally to 40 µm thickness on a vibratome.

The slices were stained with a free-floating protocol in 48-well plates. In brief, the samples were washed 4 x 15 min in PBS at RT, then permeabilized overnight in PBS + 1% (w/v) Triton-X 100 (Sigma X100-100ML). On day 2, we washed the samples 3 x 5 minutes in PBS, then blocked over the day (8 hours) in a blocking buffer (0.5% w/v Triton-X 100, 5% Neutral Horse Serum, 5% Neutral Donkey Serum, in PBS). Then, we added the primary antibodies in a fresh volume of blocking buffer (Swant anti-PV #235, mouse, 1:1000; Synaptic Systems anti-c-Fos #226 008, rabbit, 1:500). We sealed the plate with Parafilm and incubated for 48h at 4°C on a shaker. On day 5, we washed the samples 8 x 5 minutes with PBS, then incubated with the secondary antibodies in blocking buffer (Goat anti-rabbit Alexa 488, Invitrogen #A-11008, 1:500; Goat anti-mouse Alexa 568, Invitrogen A-11031, 1:500) for 3 hours at RT on a shaker in the dark. We subsequently washed the samples in PBS + 1% Triton-X 100 for 2x 15 minutes, followed by 2 x 15 minutes wash in PBS. We then counterstained the nuclei with Hoechst (Thermo Fisher # 62249) 1:5000 in PBS for 15 minutes at RT on a shaker in the dark. We then rinsed the slices in PBS and mounted them on glass slides with DAKO mounting medium (Agilent #S302380-2).

### Slice imaging and segmentation, atlas registration and region quantification

Subsequently, the slices were imaged on a Zeiss Axioskop with Apotome unit, using a Mercury lamp (Zeiss, at 58% intensity) and 5x and 10x objectives (Zeiss EC Plan-Neofluar 5 ×/0.16 M27 and EC Plan-Neofluar 10x/0.30 M27). We then stitched the images using Fiji’s collection stitching plugin with the “unknown / all files in folder” option. Where necessary, we stitched with the TrakEM plugin. The stitched images were then processed with Ilastik, using a custom-trained Random Forest model to find both c-Fos^+^ and PV^+^ neurons. The resulting probability maps were exported as 8-bit tiffs, and then further processed with a custom python script. This script thresholded the probability maps (threshold set to 128 out of 255), segmented the resulting blobs, and extracted the centerpoints as cell coordinates. Subsequently, we hand-registered every slice (400 in total) to the respective coronal image of the Allen Brain Atlas CCF3, using Fiji’s BigWarp plugin (Bogovic et al., 2016), generating a version of the atlas that was warped to the image’s contours. Our script then checked for each cell in which CCF3 atlas region it was, calculated the region’s area by counting the pixels with the same intensity (=region ID) in the slice image and calculated a per-area density cells as final result. We compared those per-region density values with unpaired t-tests that were multiple-testing corrected with 10,000 permutations. The code for this script, as well as the Ilastik weights, can be found at github https://github.com/MoritzNegwer/amygdala_paper/tree/main/Fig_4 and the European Bioimaging Archive https://doi.org/10.6019/S-BSST1975.

### Social Novelty paradigm with the LiveMouseTracker

To measure the *Ehmt1^+/−^*’s response to social novelty, we adapted a protocol from Huang and colleagues (Huang et al., 2020, 2016). We built a LiveMouseTracker (De Chaumont et al., 2019) according to the instructions in the supplementary methods. In brief, we built a stage with an overhead mounted Kinect v2 depth-sensing camera, connected to a logging PC running the LMT live-analysis software. Because we were only tracking a single mouse, the RFID chip readers at the bottom of the setup were omitted and the mice were not chipped. We built a set of four plexiglass cages (50×50×35 cm) with lids of perforated aluminum to prevent the mice from escaping by jumping over the edge. The lid was taken off only for the recording sessions. Several experimental mice – including *Ehmt1^+/−^* mice - learned to jump up against the lid and walked upside-down for several minutes (not quantified as this happened outside of the recording setup).

We filled the cage with ca. 5 cm of corncob bedding, a red plastic mouse house, food and nesting material (SizzleNest), and added a drinking water bottle through an 8 mm hole in the front plate of the cage. In the middle of the cage, we placed a 10×10×10 cm mini-cage that was empty for the duration of the experiment, until the second recording session on day 7 when it housed the stranger mouse. The mini-cage was made with walls of perforated aluminum so that the mice could hear and smell each other, but had limited exchange of tactile stimuli.

On day 1 of the experiment, we placed the empty cage in the LiveMouseTracker setup. The room was kept on a reverse day/night cycle, so that the measurements were done within the first 3 hours of the dark period. The room was dark except for the light emitted by the measurement computer screen (turned to minimum brightness). We placed a single adult female *Ehmt1^+/−^* or *Ehmt1^+/+^* mouse (termed the experimental mouse) in the 50×50×35 cm cage on day 1 of the experiment, where it remained until day 7. Mice were assigned to a cage at random at the beginning of the experiment. We measured the movement of the mouse in two subsequent sessions of 10 minutes each. Between session 1 and 2, we took out and re-placed the (empty) mini-cage to habituate the experimental mouse to the procedure. Subsequently, the lid was placed on the cage and the mouse was single-housed in its cage for the next 7 days, with one recording session per day as described above. On day 7, the first session was as before. At the start of the second session, the mini-cage was taken out as usual, and an unrelated and unknown “stranger mouse” (one of four adult wild-type C57/Bl6 females that had been group-housed in the same room to habituate to the reversed day/night cycle) was placed in the mini-cage. Once the mini-cage with the stranger mouse had been placed back, we recorded the movement of the experimental mouse around the mini-cage with the LiveMouseTracker, analyzed post-hoc with LMT software (https://github.com/fdechaumont/lmt-analysis, commit 7a1d977c9ec679f0e40c85fac834b5a493e40083 as of 2020-01-28) and custom Python scripts (https://github.com/MoritzNegwer/amygdala_paper/tree/main/Fig_5_h-k). We defined a 5 cm “contact zone” around the edge of the mini-cage and considered the experimental mouse inside this zone when the “head” coordinate was within this zone. The remaining cage was considered the “away zone”. We analyzed the time the experimental mouse spent in the away zone, and the velocity of its movements in the away zone. We also analyzed the number of sharp turns, which we used to normalize the c-Fos^+^ density data in supplementary Fig. 2. After the second session of day 7, the stranger mouse was removed from the mini-cage, the lid was placed back on the cage, and it was placed back on the shelf in the same room. The remaining 3 mice (of a total batch size of 4 mice) were measured, and exactly 2 hours after the stranger mouse exposure, the cage with the first experimental mouse was placed under a cloth cover and moved to a different room, where the experimental mouse was sacrificed under deep Isoflurane anesthesia by transcardial perfusion with ice-cold PBS followed by 4% PFA in PBS. The brains were then post-fixed in 4% PFA for 24 hours and stored in PBS with 0.05% azide until immunostainings for PV and c-Fos (Figure 4 and supplementary figure 2). The experimenter was blinded to the mouse genotype during the analysis of the data.

### Activity measurements in a social setting

Mice were genotyped around PND21 using the MyTaq Extract-PCR Kit (Meridian Bioscience, OH, USA) and subsequently weaned in test groups. Each test group consisted of two wildtype and two heterozygous mice of the same sex, that were housed together throughout the entire experiment. At PND49 (+/− 7 days) all animals were subjected to a small surgical procedure to implant a 12mm RFID chip (UNO BV, the Netherlands) for recognition in the Live Mouse Tracker (LMT) set-up. After a recovery period of one to two weeks, LMT experiments were performed using a set-up built as described by de Chaumont et al. (De Chaumont et al., 2019). During each LMT recording, a complete test group of four animals was transferred from their home cage to the LMT cage, including the shelter and nesting material, between 9.30 and 10.00am. Food and water were available ad libitum in the LMT cage. Recordings stopped the next day between 8.00 and 8.30am after which the animals were transferred back to the home cage and the set-up was thoroughly cleaned to prepare for the next recording. Experiments were recorded using the Live Mouse Tracker code in Icy and subsequently processed using the preProcessDataBase code in Icy (latest versions available on https://livemousetracker.org). Activity files were obtained by running the Compute_Distance_Lg_Term_Timebin.py script (https://github.com/fdechaumont/lmt-analysis), using 10 minutes bins and extracting the data obtained between 10.00am and 08.00am the next day to synchronize all datasets. Activity plots and statistics were performed using R Studio (2024.04.2) running R version 4.4.1. To account for large group-dependent differences in total activity, the total activity of all animals was normalized to the average total activity of the two wildtype animals in each respective group. n = 14/14 *Ehmt1^+/+^/Ehmt1^+/−^* females, and 14/14 *Ehmt1^+/+^/Ehmt1^+/−^* males.

### Image sources

The Brain slice image in Fig. 5a is from the Allen Brain Atlas (http://api.brain-map.org/api/v2/svg_download/100960033?groups=28). The mouse drawing in Fig. 5g is from NIH BioArt, licensed public domain (https://bioart.niaid.nih.gov/bioart/279), the LMT image is from the supplementary of the preprint (de Chaumont et al. Preprint at biorxiv 2018; https://www.biorxiv.org/content/10.1101/345132v2.supplementary-material), licensed CC-BY4.0 International license.

## Supporting information

Supplementary_Table_1_Friendlyclearmap_results_VIP

Supplementary_Table_2_Friendlyclearmap_results_SST

Supplementary_Table_3_Friendlyclearmap_results_PV_P14

Supplementary_Table_4_Friendlyclearmap_results_PV_P56

Supplementary_table_5_PV_P14_Amygdala_v01

Supplementary_table_6_PV_P56_Amygdala

Supplementary_Table_7_PV-cFos_Amygdala_and_frontal

Supplementary_Table_8_mIPSC_Amplitude_IEI_nested_Anova

Supplementary_Table_9_10Hz_Overview_Averages_v03

Supplementary_Table_10_LMT_behavior

## Statements & Declarations

## Funding

This work was supported by grants from the Netherlands Organization for Scientific Research, NWO-CAS grant 012.200.001 (to N.N.K) and NWO NWA.1518.22.136 SCANNER (to N.N.K.), and the Kleefstra Foundation to N.N.K, as well as a Radboudumc Internationalization Grant (for Cold Spring Harbor Asia Workshop “Building and Mining Brain Atlases and Connectomes”, Shanghai, China, 2019) and a digital travel grant for the FENS forum 2020. None of the funders had any influence on the conceptualization or execution of this study, or decision to publish.

## Competing Interests

The authors declare an absence of competing interests.

## Ethics approval

All animal experiments were conducted in conformity with the Animal Care Committee of the Radboud University Nijmegen Medical Centre and the National Committee on Animal Experiments (CCD), The Netherlands, and conform to the guidelines of the Dutch Council for Animal Care and the European Communities Council Directive 2010/63/EU (Project RU-DEC 2017-0048, permit AVD10300 2019 7746).

## Work distribution

MN and NNK conceptualized the research. MN performed the clearing, staining and electrophysiology experiments, wrote the analysis software and analyzed the data. IvdW built the LiveMouseTracker and analyzed the social activity data. AO supported the genotyping of the *Ehmt1* mice and laboratory logistics. MN, DS and NNK interpreted the data. IvdW, MN and NNK interpreted the behavioral data. NNK, HvB and DS acquired funding for this research. MN and NNK wrote the original draft of the paper, and MN, NNK, IvdW, DS and HvB provided feedback. MN, DS and NNK revised the manuscript.

## Acknowledgements

The *Ehmt1^flox/flox^* mice were a kind gift from Anne Schaefer (Rockefeller University, New York, USA; (Schaefer et al., 2009).The *PV^Cre^* and *SST^Cre^* mice were a kind gift from Tansu Celikel (now at Georgia Tech). We thank Maren Bormann, Rick Hesen, Lukas Lütje, Lynn Aarts and Anthonieke Middelman for help with the clearing experiments, and Bram Geenen, and Gert-Jan Bakker for support with the light-sheet imaging, and Bram Bosch and Carleen Rossing for support with coding the FriendlyClearMap analysis pipeline. We thank Bram Geenen, Shaghayegh Abghari, Dewi van der Geugten, Kari Bosch and Emma Clephas (all Radboudumc Nijmegen) and the members of the Ertürk lab at Helmholtz Munich for helpful discussions about tissue clearing. We would also like to thank the team of the CDL Nijmegen for support with the mouse husbandry and behavioral experiments.

## Code availability

All code specifically for this paper can be found here: https://github.com/MoritzNegwer/amygdala_paper under a GLP v3 license

The code for FriendlyClearMap can be found here: https://github.com/MoritzNegwer/FriendlyClearMap-scripts under a GPL v3 license (see also Negwer et al. Gigascience 2023)

## Data availability

All data for this publication can be found on BioStudies / European Bioimage Archive under a CC0 (Public Domain) license:

- https://doi.org/10.6019/S-BSST1975

Supplementary tables 1-10 are available at EBI under a CC0 (Public Domain) license:

- https://ftp.ebi.ac.uk/pub/databases/biostudies/S-BSST/975/S-BSST1975/Files/ Supplementary_Tables

Supplementary videos are available at EBI under a CC0 (Public Domain) license:

- https://ftp.ebi.ac.uk/pub/databases/biostudies/S-BSST/975/S-BSST1975/Files/ Supp_video_01_VIP_WT.mp4
- https://ftp.ebi.ac.uk/pub/databases/biostudies/S-BSST/975/S-BSST1975/Files/ Supp_video_02_SST_WT.mp4
- https://ftp.ebi.ac.uk/pub/databases/biostudies/S-BSST/975/S-BSST1975/Files/ Supp_video_03_PV-P14_WT.mp4
- https://ftp.ebi.ac.uk/pub/databases/biostudies/S-BSST/975/S-BSST1975/Files/ Supp_video_04_PV-P56_WT.mp4

---

**Supplementary Figure 1, related to Figure 2.**
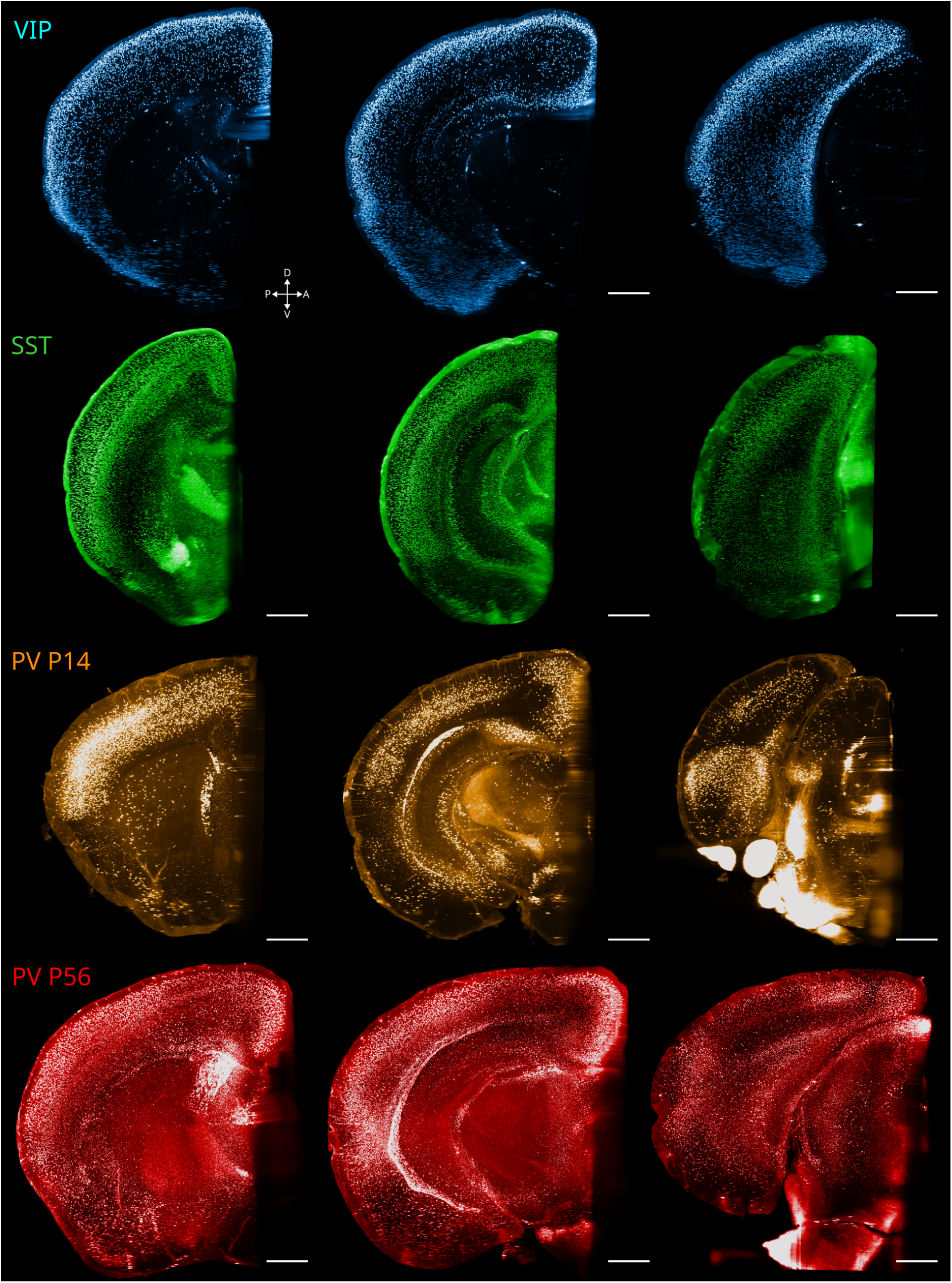
Representative coronal planes of the four reporter strains we used, to show that the anti-tdTomato antibodies penetrated evenly throughout the hemisphere. VIP^+^ neurons are almost exclusively found in the cortical plate, which is why the staining appears as a shell. For the SST^+^ hemispheres, we noticed a bright staining on the outside of the cortex (though not the non-cortical surfaces, e.g. midline cutting plane), which we interpreted as brightly stained layer-1 projecting axons. Nonetheless, the inside of the hemisphere was evenly stained. For PV-P14 and PV-P56, the brightest staining was visible in the cerebellum, which we ommitted during our analysis due to the extreme brightness that saturated our microscope’s detector. Nonetheless, even dense structures inside the hemishpere such as the thalamic reticular nucleus were stained evenly. Some of those images have been previously used in Negwer et al., 2022; reused here under Creative Commons Attribution (CC-BY) 4.0 license.

**Supplementary Figure 2, related to Figure 4.**
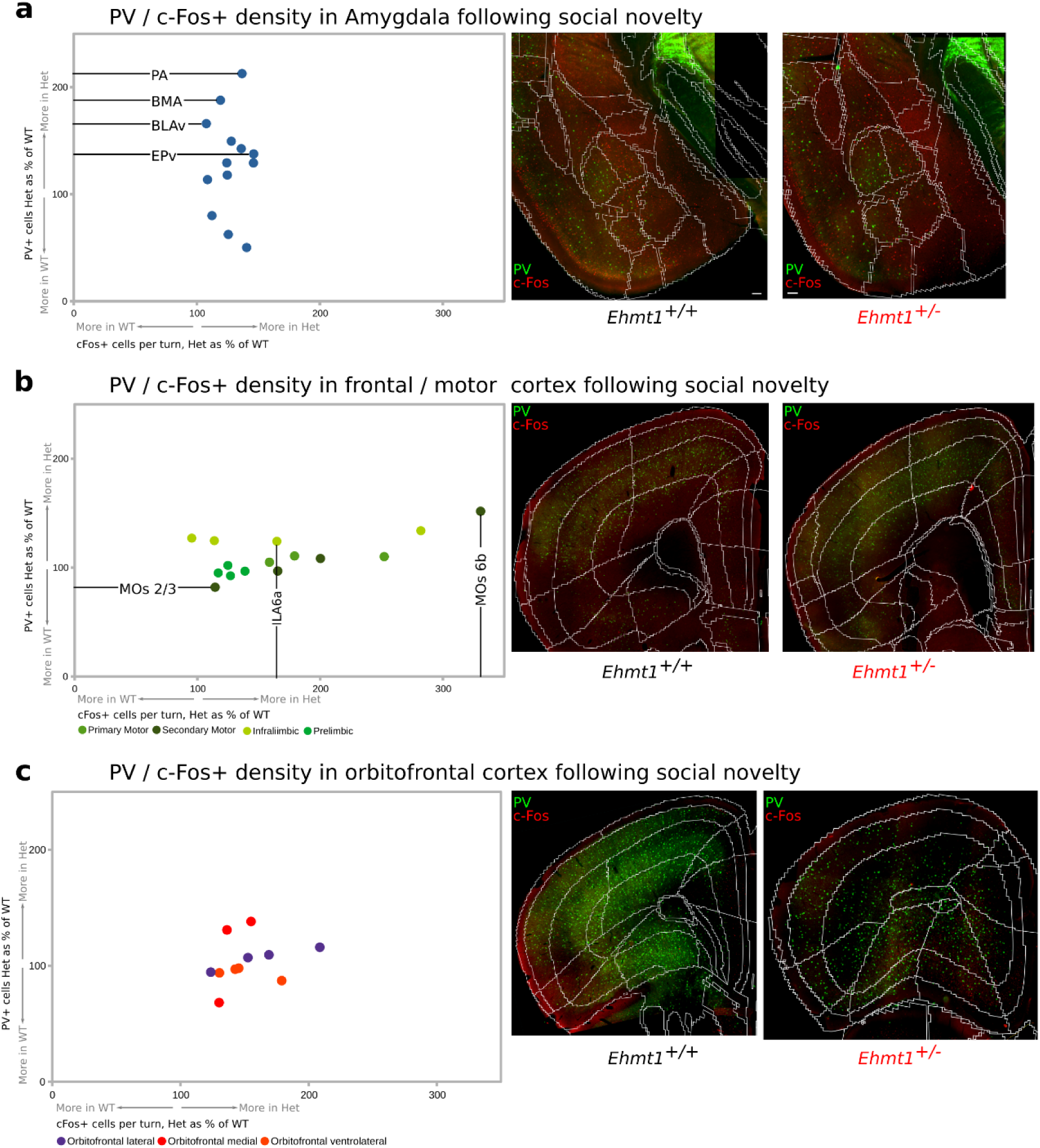
c-Fos^+^ and PV^+^ cell density following social novelty. Y-axis: Relative PV^+^ density (*Ehmt1^+/−^* normalized to *Ehmt1^+/+^*). X-axis: Relative c-Fos^+^ cell density (*Ehmt1^+/−^* normalized to *Ehmt1^+/+^*) per area, normalized to the mouse’s activity (number of sharp turns) during the social novelty challenge. The number of turns was unchanged between genotypes during the social novelty challenge, see Fig. 5 k. Statistically significant differences are indicated with a horizontal (for PV^+^ density) or horizontal (for movement-normalized c-Fos^+^ density) line and the abbreviation of the region. **a)** in the amygdala, the density of PV^+^ neurons is significantly higher in the *Ehmt1^+/−^* in the PA, BMA, BLAv and EPv regions (this is the same data as Fig. 4 c). In contrast, the (movement-normalized) density of c-Fos^+^ neurons is unchanged between genotypes. Right side: Amygdala slices of *Ehmt1^+/+^* (left) and *Ehmt1^+/−^* (right), with the area borders superimposed, immunostained against c-Fos (red) and PV (green), those mice do not have a tdTomato reporter. **b)** in the motor cortex (primary motor cortex: light green, secondary motor cortex: darker green), prelimbic cortex (yellow green) and infralimbic cortex (bright green), the PV^+^ density is only higher in layers 2/3 of the secondary motor cortex (MOs2/3, horizontal line). In contrast, the motor cortex fields show a change in (movement-normalized) c-Fos^+^ density between genotypes, which is significantly higher in the *Ehmt1^+/−^* infralimbic layer 6a (IL6a) and secondary motor cortex layer 6b (MOs6b). This indicates that the *Ehmt1^+/−^* motor cortex requires more activation for the same amount of movement. **c)** In the orbitofrontal cortex, there is no change in either (movement-normalized) c-Fos^+^ density or PV^+^ density between genotypes. Each dot: Average of one region across 48/48 slices from 12/12 *Ehmt1^+/+^ / Ehmt1^+/−^* mice, all female. Please see supplementary table 7 for details.

**Supplementary Figure 3, related to Figure 5.**
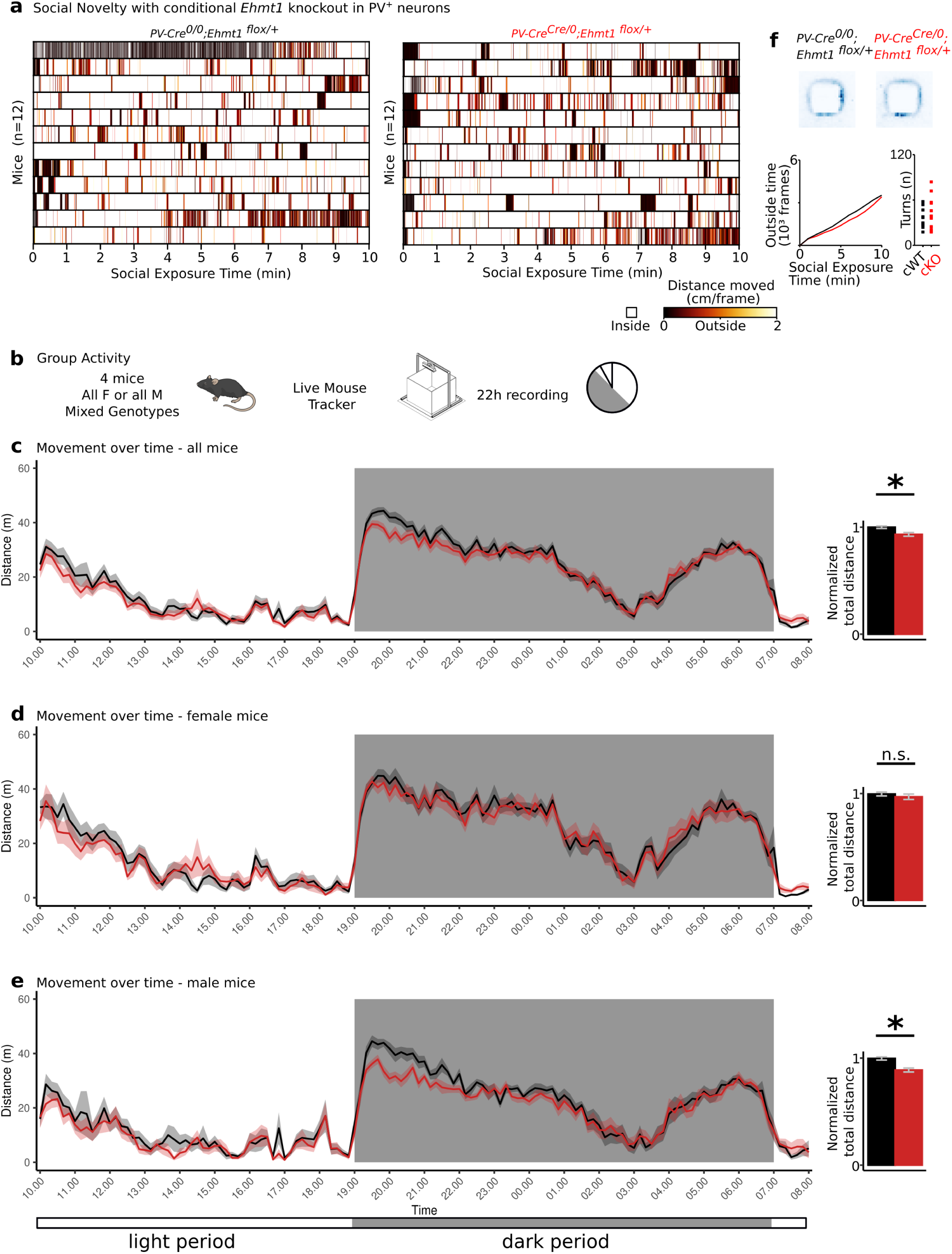
**a)** Reaction to social novelty challenge is unchanged in conditional *Ehmt1* knockout in PV^+^ neurons. Neither the movement patterns (left), time spent in the outside zone (right), or number of turns (bar chart on the right) is changed between genotypes. N= 12/12 *PV^Cre^;Ehmt1^+/+^* / *PV^Cre^;Ehmt1^flox/+^*, all females. Please see supplementary table 10 for details. **b)** 22h-activity shows nighttime hypoactivity in males a social context. Mice were placed in a LiveMouseTracker in groups of 4 (same-sex cage mates, mixed genotypes) and tracked for 22 hours. **c)** Combined activity patterns of male and female mice show an activity reduction in *Ehmt1^+/−^* mice at the onset of the dark period (20:00 – 22:00). **d)** This difference is not evident in female mice, where activity levels between *Ehmt1^+/+^* and *Ehmt1^+/−^* mice are indistinguishable. **e)** instead, the difference in activity levels is driven by male mice, where there is a significant reduction in activity levels at the onset of the dark period in the *Ehmt1^+/−^* males. N = 14/14 *Ehmt1^+/+^/Ehmt1^+/−^* females, and 14/14 *Ehmt1^+/+^/Ehmt1^+/−^* males. p (all mice) = 0.00213 (two-sided t-test), p (females) = 0.4102 (Wilcoxon test because samples were not normally distributed), p (males) = 0.0001102 (two-sided t-test).

**Supplementary Figure 4:**
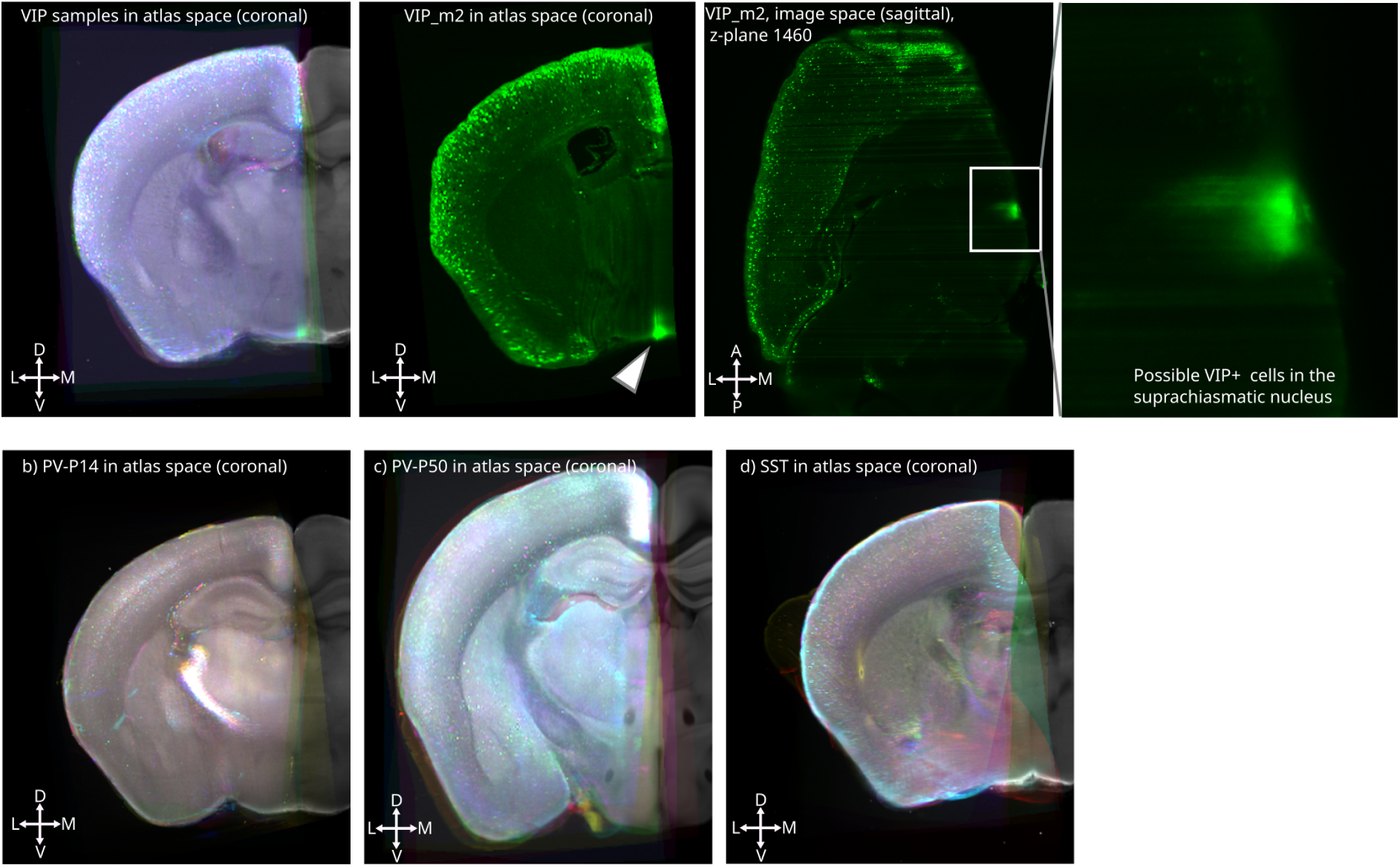
Overlap of all slices in atlas space. **a)** VIP samples in atlas space (overlay, one color per brain. The atlas is in gray, visible to the right of the first image. Not all samples are cut exactly the same along the midline, which means that e.g. the suprachiasmatic nucleus (arrowhead) is not evenly captured in all samples. **b)** PV-P14 atlas overlap, **c)** PV-P50 atlas overlap, **d)** SST atlas overlap.

**Supplementary Figure 5:**
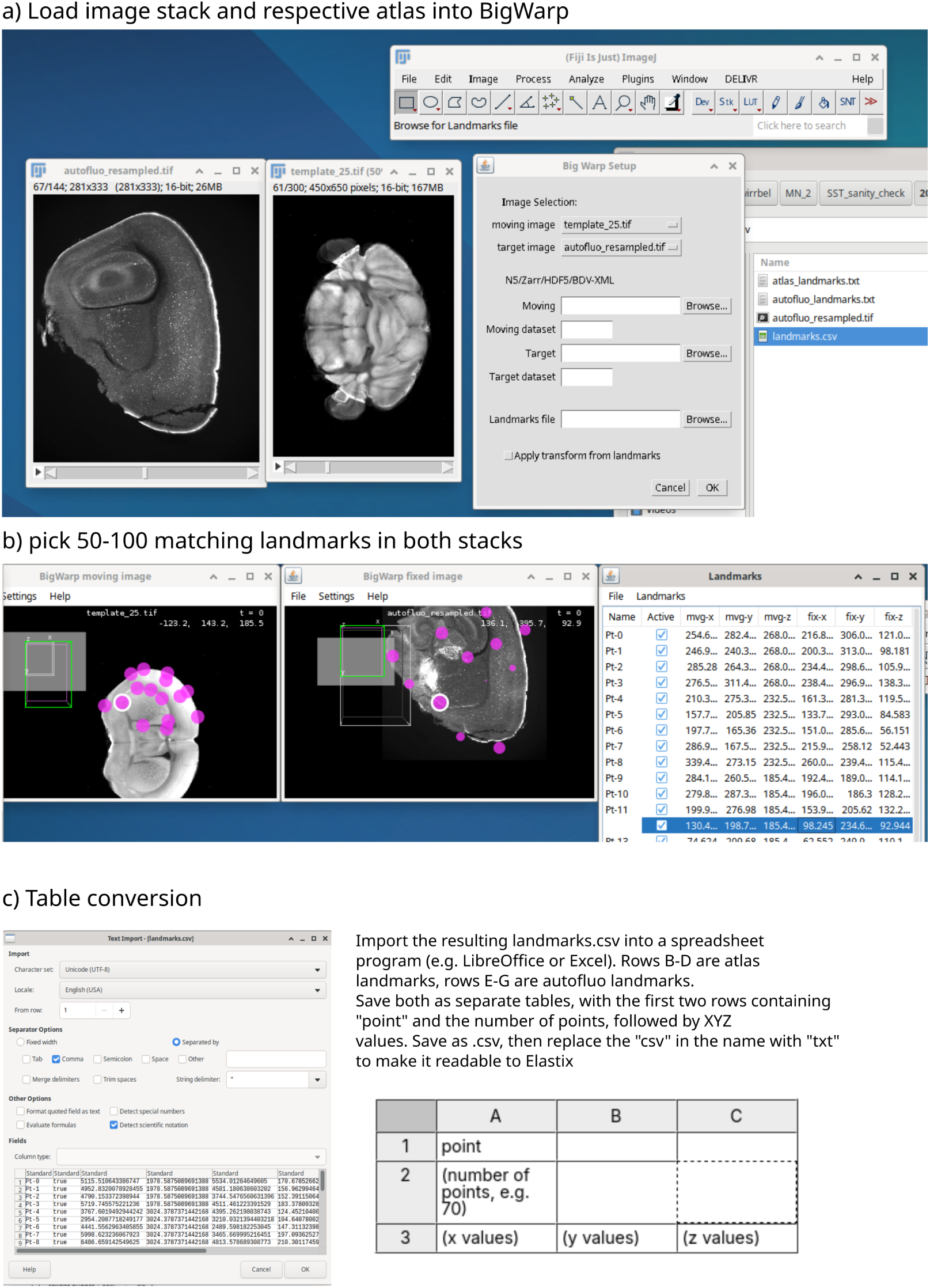
Landmark selection procedure. **a)** Loading the image stack and respective atlas file into BigWarp. **b)** Select 50-100 landmarks between the downsampled brain stack and the respective atlas stack. We paid special attention to the following regions for landmark selection: Hippocampus (CA1, DG), Cortex (pia and white matter in an imagined column in e.g. S1 barrel field and the frontal pole of the PFC, visual cortex, motor cortex and retrosplenial cortex), thalamic reticular nucleus, fiber bundles, hypothalamus, amygdala. **c)** Manual conversion of the landmarks into separate atlas_landmarks.txt and brain_landmarks.txt files.

**Supplementary Figure 6:**
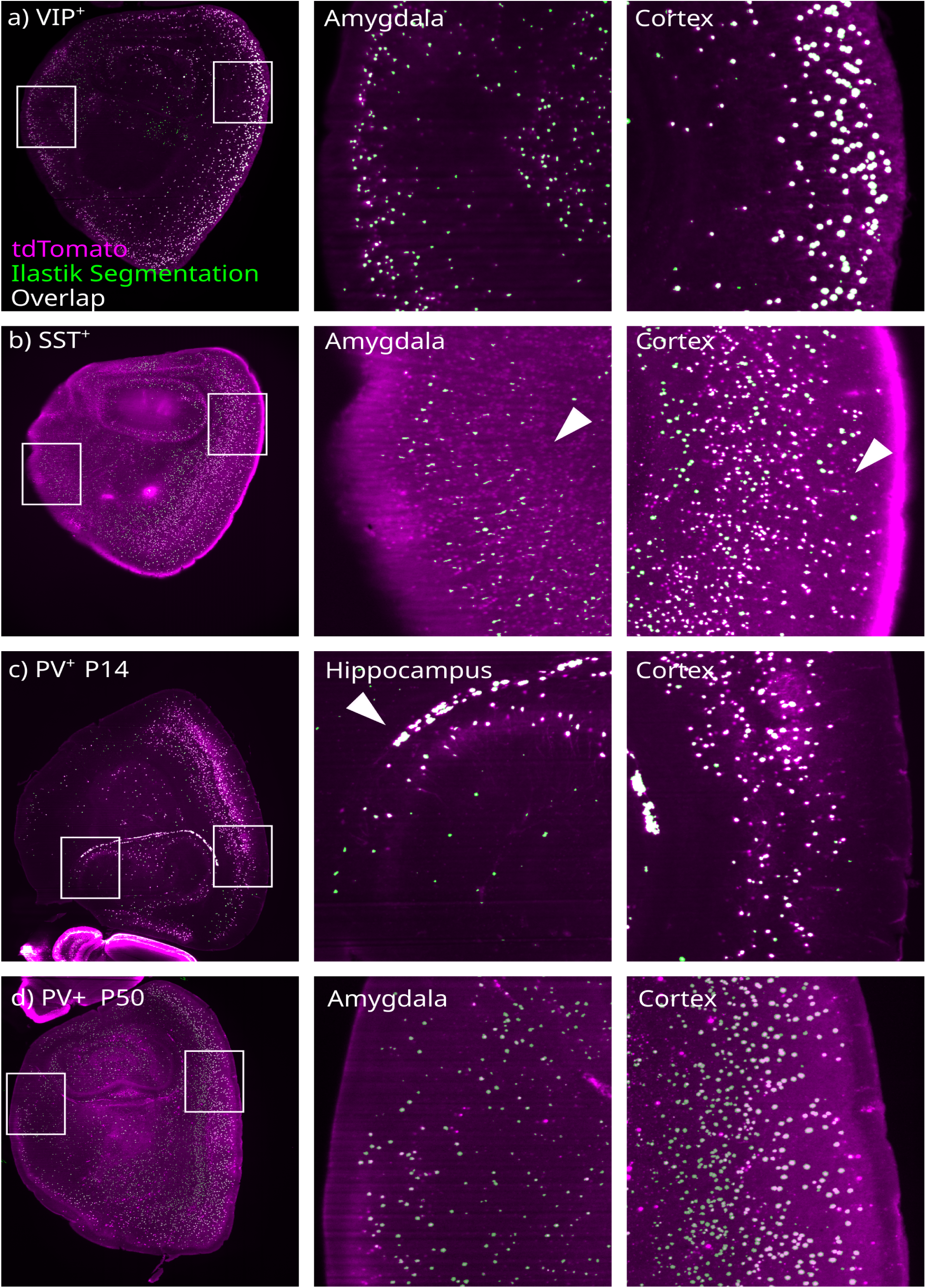
Segmentation performance of FriendlyClearMap. Overlay of anti-tdTomato signal (magenta) and Ilastik’s segmentation results (green), overlapping (true positives) in white. **a)** VIP-tdTom, sagittal overview and close-up of amygdala and and cortex. Note the comparably low anti-tdTomato signal in the amygdala that was still correctly detected as cells. **b)** SST-tdTom, sagittal overview and amygdala and cortex close-ups. Note the missed cells (false negatives, arrowheads) in the amygdala and partially in the cortex. **c)** PV P14-tdTom, sagittal overview and hippocampus and cortex close-ups. Most cells were correctly recognized, but so were some bright off-target regions along the hippocampal edge (arrowhead), presumably tdTom^+^ axons in the fiber tracts. **d)** PV P50-tdTom, sagittal overview and amygdala and cortical close-up. Note that despite brightness differences in the anti-tdTomato channel, cells are still correctly recognized.

**Supplementary Figure 7:**
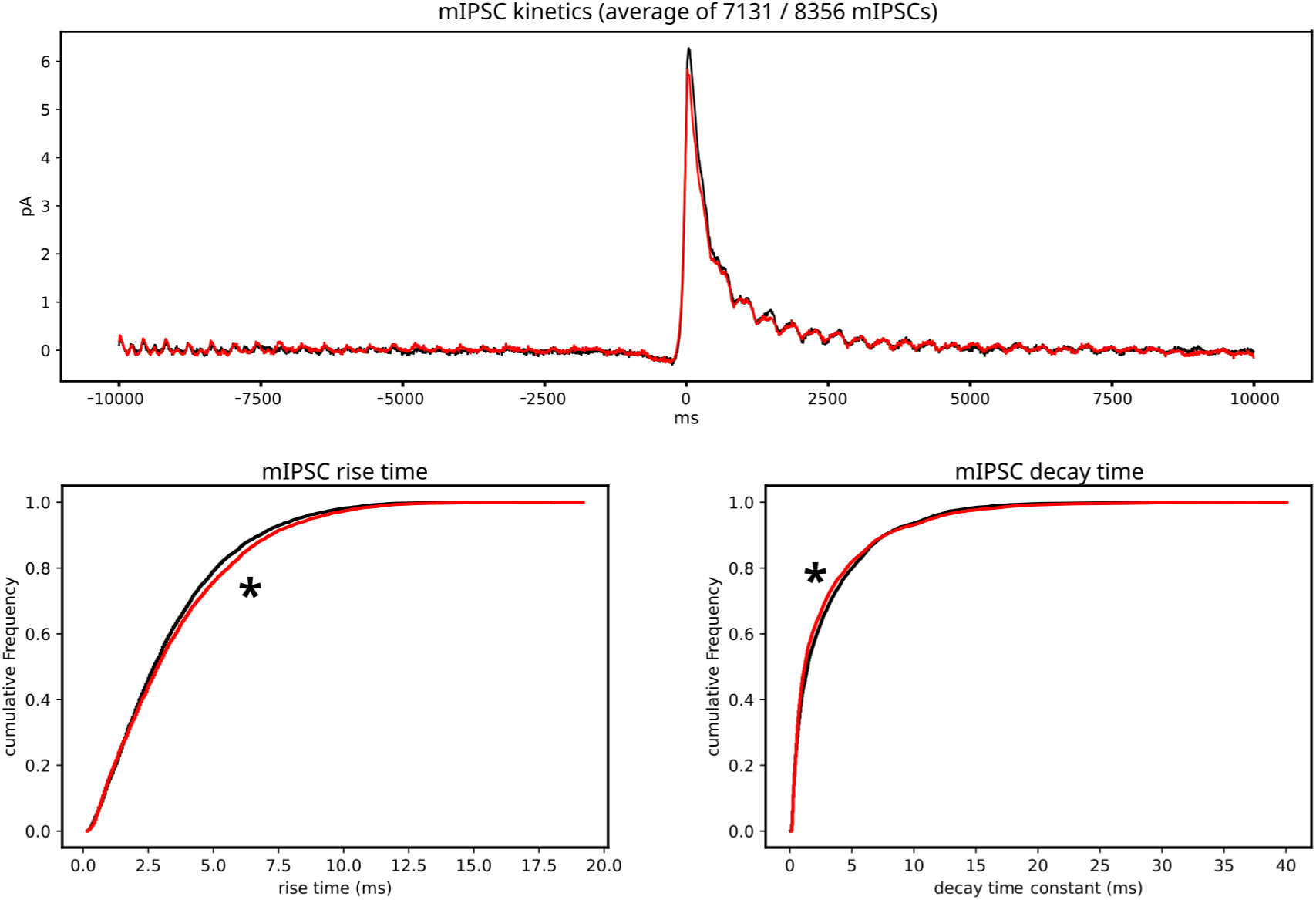
mIPSC kinetics. Taken across all mIPSCs (n = 7131/8356 for *Ehmt1^+/+^*/*Ehmt1^+/−^*). Top row: Average mIPSC time course, centered around the onset. Bottom row: Cumulative plots of mIPSC rise and decay time. * = significant, 2-sample Kolmogorov-Smirnov test (rise time p = 0.0001508606; decay time p = 0.0000000007).

## Notes

https://doi.org/10.6019/S-BSST1975

https://github.com/MoritzNegwer/amygdala_paper

https://github.com/MoritzNegwer/FriendlyClearMap-scripts

### Competing Interest Statement

The authors have declared no competing interest.

### Summary of Updates

New figure 2c and new supplementary figures 1, 4, 5, 6 and 7, refinements throughout the text. Supplementary tables 1-4 updated.

## References

Achim, K., 2024. Epigenetic and transcriptional regulation of neuron phenotype. Int. J. Dev. Biol. 68, 199–209. 10.1387/ijdb.230204ka

Aerts, T., Seuntjens, E., 2021. Novel Perspectives on the Development of the Amygdala in Rodents. Front. Neuroanat. 15, 786679. 10.3389/fnana.2021.786679

Allaway, K.C., Gabitto, M.I., Wapinski, O., Saldi, G., Wang, C.-Y., Bandler, R.C., Wu, S.J., Bonneau, R., Fishell, G., 2021. Genetic and epigenetic coordination of cortical interneuron development. Nature 597, 693–697. 10.1038/s41586-021-03933-1

Apicella, A.J., Marchionni, I., 2022. VIP-Expressing GABAergic Neurons: Disinhibitory vs. Inhibitory Motif and Its Role in Communication Across Neocortical Areas. Front. Cell. Neurosci. 16, 811484. 10.3389/fncel.2022.811484

Bakken, T.E., Jorstad, N.L., Hu, Q., Lake, B.B., Tian, W., Kalmbach, B.E., Crow, M., Hodge, R.D., Krienen, F.M., Sorensen, S.A., Eggermont, J., Yao, Z., Aevermann, B.D., Aldridge, A.I., Bartlett, A., Bertagnolli, D., Casper, T., Castanon, R.G., Crichton, K., Daigle, T.L., Dalley, R., Dee, N., Dembrow, N., Diep, D., Ding, S.-L., Dong, W., Fang, R., Fischer, S., Goldman, M., Goldy, J., Graybuck, L.T., Herb, B.R., Hou, X., Kancherla, J., Kroll, M., Lathia, K., Van Lew, B., Li, Y.E., Liu, C.S., Liu, H., Lucero, J.D., Mahurkar, A., McMillen, D., Miller, J.A., Moussa, M., Nery, J.R., Nicovich, P.R., Niu, S.-Y., Orvis, J., Osteen, J.K., Owen, S., Palmer, C.R., Pham, T., Plongthongkum, N., Poirion, O., Reed, N.M., Rimorin, C., Rivkin, A., Romanow, W.J., Sedeño-Cortés, A.E., Siletti, K., Somasundaram, S., Sulc, J., Tieu, M., Torkelson, A., Tung, H., Wang, X., Xie, F., Yanny, A.M., Zhang, R., Ament, S.A., Behrens, M.M., Bravo, H.C., Chun, J., Dobin, A., Gillis, J., Hertzano, R., Hof, P.R., Höllt, T., Horwitz, G.D., Keene, C.D., Kharchenko, P.V., Ko, A.L., Lelieveldt, B.P., Luo, C., Mukamel, E.A., Pinto-Duarte, A., Preissl, S., Regev, A., Ren, B., Scheuermann, R.H., Smith, K., Spain, W.J., White, O.R., Koch, C., Hawrylycz, M., Tasic, B., Macosko, E.Z., McCarroll, S.A., Ting, J.T., Zeng, H., Zhang, K., Feng, G., Ecker, J.R., Linnarsson, S., Lein, E.S., 2021. Comparative cellular analysis of motor cortex in human, marmoset and mouse. Nature 598, 111–119. 10.1038/s41586-021-03465-8

Bakker, R., Tiesinga, P., Kötter, R., 2015. The Scalable Brain Atlas: Instant Web-Based Access to Public Brain Atlases and Related Content. Neuroinform 13, 353–366. 10.1007/s12021-014-9258-x

Balemans, M.C.M., Ansar, M., Oudakker, A.R., Van Caam, A.P.M., Bakker, B., Vitters, E.L., Van Der Kraan, P.M., De Bruijn, D.R.H., Janssen, S.M., Kuipers, A.J., Huibers, M.M.H., Maliepaard, E.M., Walboomers, X.F., Benevento, M., Nadif Kasri, N., Kleefstra, T., Zhou, H., Van Der Zee, C.E.E.M., Van Bokhoven, H., 2014. Reduced Euchromatin histone methyltransferase 1 causes developmental delay, hypotonia, and cranial abnormalities associated with increased bone gene expression in Kleefstra syndrome mice. Developmental Biology 386, 395–407. 10.1016/j.ydbio.2013.12.016

Balemans, M.C.M., Huibers, M.M.H., Eikelenboom, N.W.D., Kuipers, A.J., Van Summeren, R.C.J., Pijpers, M.M.C.A., Tachibana, M., Shinkai, Y., Van Bokhoven, H., Van Der Zee, C.E.E.M., 2010. Reduced exploration, increased anxiety, and altered social behavior: Autistic-like features of euchromatin histone methyltransferase 1 heterozygous knockout mice. Behavioural Brain Research 208, 47–55. 10.1016/j.bbr.2009.11.008

Balemans, M.C.M., Nadif Kasri, N., Kopanitsa, M.V., Afinowi, N.O., Ramakers, G., Peters, T.A., Beynon, A.J., Janssen, S.M., Van Summeren, R.C.J., Eeftens, J.M., Eikelenboom, N., Benevento, M., Tachibana, M., Shinkai, Y., Kleefstra, T., Van Bokhoven, H., Van Der Zee, C.E.E.M., 2013. Hippocampal dysfunction in the Euchromatin histone methyltransferase 1 heterozygous knockout mouse model for Kleefstra syndrome. Human Molecular Genetics 22, 852–866. 10.1093/hmg/dds490

Benevento, M., Iacono, G., Selten, M., Ba, W., Oudakker, A., Frega, M., Keller, J., Mancini, R., Lewerissa, E., Kleefstra, T., Stunnenberg, H.G., Zhou, H., Van Bokhoven, H., Nadif Kasri, N., 2016. Histone Methylation by the Kleefstra Syndrome Protein EHMT1 Mediates Homeostatic Synaptic Scaling. Neuron 91, 341–355. 10.1016/j.neuron.2016.06.003

Benevento, M., Oomen, C.A., Horner, A.E., Amiri, H., Jacobs, T., Pauwels, C., Frega, M., Kleefstra, T., Kopanitsa, M.V., Grant, S.G.N., Bussey, T.J., Saksida, L.M., Van Der Zee, C.E.E.M., Van Bokhoven, H., Glennon, J.C., Kasri, N.N., 2017. Haploinsufficiency of EHMT1 improves pattern separation and increases hippocampal cell proliferation. Sci Rep 7, 40284. 10.1038/srep40284

Benevento, M., Van De Molengraft, M., Van Westen, R., Van Bokhoven, H., Nadif Kasri, N., 2015. The role of chromatin repressive marks in cognition and disease: A focus on the repressive complex GLP/G9a. Neurobiology of Learning and Memory 124, 88–96. 10.1016/j.nlm.2015.06.013

Berg, S., Kutra, D., Kroeger, T., Straehle, C.N., Kausler, B.X., Haubold, C., Schiegg, M., Ales, J., Beier, T., Rudy, M., Eren, K., Cervantes, J.I., Xu, B., Beuttenmueller, F., Wolny, A., Zhang, C., Koethe, U., Hamprecht, F.A., Kreshuk, A., 2019. ilastik: interactive machine learning for (bio)image analysis. Nat Methods 16, 1226–1232. 10.1038/s41592-019-0582-9

Bogovic, J.A., Hanslovsky, P., Wong, A., Saalfeld, S., 2016. Robust registration of calcium images by learned contrast synthesis, in: 2016 IEEE 13th International Symposium on Biomedical Imaging (ISBI). Presented at the 2016 IEEE 13th International Symposium on Biomedical Imaging (ISBI 2016), IEEE, Prague, Czech Republic, pp. 1123–1126. 10.1109/ISBI.2016.7493463

Cai, R., Pan, C., Ghasemigharagoz, A., Todorov, M.I., Foerstera, B., Zhao, S., Bhatia, H.S., Mrowka, L., Theodorou, D., Rempfler, M., Xavier, A., Kress, B.T., Benakis, C., Liesz, A., Menze, B., Kerschensteiner, M., Nedergaard, M., Erturk, A., 2018. Panoptic vDISCO imaging reveals neuronal connectivity, remote trauma effects and meningeal vessels in intact transparent mice. 10.1101/374785

Cattani, A., Arnold, D.B., McCarthy, M., Kopell, N., 2024. Basolateral amygdala oscillations enable fear learning in a biophysical model. eLife 12, RP89519. 10.7554/eLife.89519

Courtin, J., Karalis, N., Gonzalez-Campo, C., Wurtz, H., Herry, C., 2014. Persistence of amygdala gamma oscillations during extinction learning predicts spontaneous fear recovery. Neurobiology of Learning and Memory 113, 82–89. 10.1016/j.nlm.2013.09.015

De Chaumont, F., Ey, E., Torquet, N., Lagache, T., Dallongeville, S., Imbert, A., Legou, T., Le Sourd, A.-M., Faure, P., Bourgeron, T., Olivo-Marin, J.-C., 2019. Real-time analysis of the behaviour of groups of mice via a depth-sensing camera and machine learning. Nat Biomed Eng 3, 930–942. 10.1038/s41551-019-0396-1

Del Rio, J., De Lecea, L., Ferrer, I., Soriano, E., 1994. The development of parvalbumin-immunoreactivity in the neocortex of the mouse. Developmental Brain Research 81, 247–259. 10.1016/0165-3806(94)90311-5

Dervinis, M., Major, G., 2022. A novel method for reliably measuring miniature and spontaneous postsynaptic potentials/currents in whole-cell patch clamp recordings in the central nervous system. 10.1101/2022.03.20.485046

Elmqvist, D., Quastel, D.M., 1965. A quantitative study of end-plate potentials in isolated human muscle. The Journal of Physiology 178, 505–529. 10.1113/jphysiol.1965.sp007639

Fagiolini, M., Hensch, T.K., 2000. Inhibitory threshold for critical-period activation in primary visual cortex. Nature 404, 183–186. 10.1038/35004582

Favuzzi, E., Deogracias, R., Marques-Smith, A., Maeso, P., Jezequel, J., Exposito-Alonso, D., Balia, M., Kroon, T., Hinojosa, A.J., F. Maraver, E., Rico, B., 2019. Distinct molecular programs regulate synapse specificity in cortical inhibitory circuits. Science 363, 413–417. 10.1126/science.aau8977

Feng, F., Headley, D.B., Amir, A., Kanta, V., Chen, Z., Paré, D., Nair, S.S., 2019. Gamma Oscillations in the Basolateral Amygdala: Biophysical Mechanisms and Computational Consequences. eNeuro 6, ENEURO.0388-18.2018. 10.1523/ENEURO.0388-18.2018

Fishell, G., Kepecs, A., 2020. Interneuron Types as Attractors and Controllers. Annu. Rev. Neurosci. 43, 1–30. 10.1146/annurev-neuro-070918-050421

Fisher, J., Verhagen, M., Long, Z., Moissidis, M., Yan, Y., He, C., Wang, J., Micoli, E., Alastruey, C.M., Moors, R., Marín, O., Mi, D., Lim, L., 2024. Cortical somatostatin long-range projection neurons and interneurons exhibit divergent developmental trajectories. Neuron 112, 558–573.e8. 10.1016/j.neuron.2023.11.013

Fogarty, M., Grist, M., Gelman, D., Marín, O., Pachnis, V., Kessaris, N., 2007. Spatial Genetic Patterning of the Embryonic Neuroepithelium Generates GABAergic Interneuron Diversity in the Adult Cortex. J. Neurosci. 27, 10935–10946. 10.1523/JNEUROSCI.1629-07.2007

Frega, M., Linda, K., Keller, J.M., Gümüş-Akay, G., Mossink, B., Van Rhijn, J.-R., Negwer, M., Gunnewiek, T.K., Foreman, K., Kompier, N., Schoenmaker, C., Van Den Akker, W., Oudakker, A., Zhou, H., Kleefstra, T., Schubert, D., Van Bokhoven, H., Kasri, N.N., 2019. Neuronal network dysfunction in a human model for Kleefstra syndrome mediated by enhanced NMDAR signaling. 10.1101/585596

Gouwens, N.W., Sorensen, S.A., Baftizadeh, F., Budzillo, A., Lee, B.R., Jarsky, T., Alfiler, L., Baker, K., Barkan, E., Berry, K., Bertagnolli, D., Bickley, K., Bomben, J., Braun, T., Brouner, K., Casper, T., Crichton, K., Daigle, T.L., Dalley, R., De Frates, R.A., Dee, N., Desta, T., Lee, S.D., Dotson, N., Egdorf, T., Ellingwood, L., Enstrom, R., Esposito, L., Farrell, C., Feng, D., Fong, O., Gala, R., Gamlin, C., Gary, A., Glandon, A., Goldy, J., Gorham, M., Graybuck, L., Gu, H., Hadley, K., Hawrylycz, M.J., Henry, A.M., Hill, D., Hupp, M., Kebede, S., Kim, T.K., Kim, L., Kroll, M., Lee, C., Link, K.E., Mallory, M., Mann, R., Maxwell, M., McGraw, M., McMillen, D., Mukora, A., Ng, Lindsay, Ng, Lydia, Ngo, K., Nicovich, P.R., Oldre, A., Park, D., Peng, H., Penn, O., Pham, T., Pom, A., Popović, Z., Potekhina, L., Rajanbabu, R., Ransford, S., Reid, D., Rimorin, C., Robertson, M., Ronellenfitch, K., Ruiz, A., Sandman, D., Smith, K., Sulc, J., Sunkin, S.M., Szafer, A., Tieu, M., Torkelson, A., Trinh, J., Tung, H., Wakeman, W., Ward, K., Williams, G., Zhou, Z., Ting, J.T., Arkhipov, A., Sümbül, U., Lein, E.S., Koch, C., Yao, Z., Tasic, B., Berg, J., Murphy, G.J., Zeng, H., 2020. Integrated Morphoelectric and Transcriptomic Classification of Cortical GABAergic Cells. Cell 183, 935–953.e19. 10.1016/j.cell.2020.09.057

Gouwens, N.W., Sorensen, S.A., Berg, J., Lee, C., Jarsky, T., Ting, J., Sunkin, S.M., Feng, D., Anastassiou, C.A., Barkan, E., Bickley, K., Blesie, N., Braun, T., Brouner, K., Budzillo, A., Caldejon, S., Casper, T., Castelli, D., Chong, P., Crichton, K., Cuhaciyan, C., Daigle, T.L., Dalley, R., Dee, N., Desta, T., Ding, S.-L., Dingman, S., Doperalski, A., Dotson, N., Egdorf, T., Fisher, M., De Frates, R.A., Garren, E., Garwood, M., Gary, A., Gaudreault, N., Godfrey, K., Gorham, M., Gu, H., Habel, C., Hadley, K., Harrington, J., Harris, J.A., Henry, A., Hill, D., Josephsen, S., Kebede, S., Kim, L., Kroll, M., Lee, B., Lemon, T., Link, K.E., Liu, X., Long, B., Mann, R., McGraw, M., Mihalas, S., Mukora, A., Murphy, G.J., Ng, Lindsay, Ngo, K., Nguyen, T.N., Nicovich, P.R., Oldre, A., Park, D., Parry, S., Perkins, J., Potekhina, L., Reid, D., Robertson, M., Sandman, D., Schroedter, M., Slaughterbeck, C., Soler-Llavina, G., Sulc, J., Szafer, A., Tasic, B., Taskin, N., Teeter, C., Thatra, N., Tung, H., Wakeman, W., Williams, G., Young, R., Zhou, Z., Farrell, C., Peng, H., Hawrylycz, M.J., Lein, E., Ng, Lydia, Arkhipov, A., Bernard, A., Phillips, J.W., Zeng, H., Koch, C., 2019. Classification of electrophysiological and morphological neuron types in the mouse visual cortex. Nat Neurosci 22, 1182–1195. 10.1038/s41593-019-0417-0

Hájos, N., 2021. Interneuron Types and Their Circuits in the Basolateral Amygdala. Front. Neural Circuits 15, 687257. 10.3389/fncir.2021.687257

Hintiryan, H., Bowman, I., Johnson, D.L., Korobkova, L., Zhu, M., Khanjani, N., Gou, L., Gao, L., Yamashita, S., Bienkowski, M.S., Garcia, L., Foster, N.N., Benavidez, N.L., Song, M.Y., Lo, D., Cotter, K.R., Becerra, M., Aquino, S., Cao, C., Cabeen, R.P., Stanis, J., Fayzullina, M., Ustrell, S.A., Boesen, T., Tugangui, A.J., Zhang, Z.-G., Peng, B., Fanselow, M.S., Golshani, P., Hahn, J.D., Wickersham, I.R., Ascoli, G.A., Zhang, L.I., Dong, H.-W., 2021. Connectivity characterization of the mouse basolateral amygdalar complex. Nat Commun 12, 2859. 10.1038/s41467-021-22915-5

Hintiryan, H., Dong, H.-W., 2022. Brain Networks of Connectionally Unique Basolateral Amygdala Cell Types. J Exp Neurosci 17, 26331055221080175. 10.1177/26331055221080175

Hochgerner, H., Singh, S., Tibi, M., Lin, Z., Skarbianskis, N., Admati, I., Ophir, O., Reinhardt, N., Netser, S., Wagner, S., Zeisel, A., 2023. Neuronal types in the mouse amygdala and their transcriptional response to fear conditioning. Nat Neurosci 26, 2237–2249. 10.1038/s41593-023-01469-3

Huang, W.-C., Chen, Y., Page, D.T., 2016. Hyperconnectivity of prefrontal cortex to amygdala projections in a mouse model of macrocephaly/autism syndrome. Nat Commun 7, 13421. 10.1038/ncomms13421

Huang, W.-C., Zucca, A., Levy, J., Page, D.T., 2020. Social Behavior Is Modulated by Valence-Encoding mPFC-Amygdala Sub-circuitry. Cell Reports 32, 107899. 10.1016/j.celrep.2020.107899

Huang, Z.J., Paul, A., 2019. The diversity of GABAergic neurons and neural communication elements. Nat Rev Neurosci 20, 563–572. 10.1038/s41583-019-0195-4

Iacono, G., Dubos, A., Méziane, H., Benevento, M., Habibi, E., Mandoli, A., Riet, F., Selloum, M., Feil, R., Zhou, H., Kleefstra, T., Kasri, N.N., van Bokhoven, H., Herault, Y., Stunnenberg, H.G., 2018. Increased H3K9 methylation and impaired expression of Protocadherins are associated with the cognitive dysfunctions of the Kleefstra syndrome. Nucleic Acids Res 46, 4950–4965. 10.1093/nar/gky196

Jackson, A.D., Cohen, J.L., Phensy, A.J., Chang, E.F., Dawes, H.E., Sohal, V.S., 2024. Amygdala-hippocampus somatostatin interneuron beta-synchrony underlies a cross-species biomarker of emotional state. Neuron 112, 1182–1195.e5. 10.1016/j.neuron.2023.12.017

Kahan, A., Mahe, K., Dutta, S., Kassraian, P., Wang, A., Gradinaru, V., 2023. Immediate responses to ambient light in vivo reveal distinct subpopulations of suprachiasmatic VIP neurons. iScience 26, 107865. 10.1016/j.isci.2023.107865

Kawaguchi, Y., Kondo, S., 2002. Parvalbumin, somatostatin and cholecystokinin as chemical markers for specific GABAergic interneuron types in the rat frontal cortex. J Neurocytol 31, 277–287. 10.1023/A:1024126110356

Kessaris, N., Denaxa, M., 2023. Cortical interneuron specification and diversification in the era of big data. Current Opinion in Neurobiology 80, 102703. 10.1016/j.conb.2023.102703

Kessaris, N., Magno, L., Rubin, A.N., Oliveira, M.G., 2014. Genetic programs controlling cortical interneuron fate. Current Opinion in Neurobiology 26, 79–87. 10.1016/j.conb.2013.12.012

Kim, Y., Yang, G.R., Pradhan, K., Venkataraju, K.U., Bota, M., García Del Molino, L.C., Fitzgerald, G., Ram, K., He, M., Levine, J.M., Mitra, P., Huang, Z.J., Wang, X.-J., Osten, P., 2017. Brain-wide Maps Reveal Stereotyped Cell-Type-Based Cortical Architecture and Subcortical Sexual Dimorphism. Cell 171, 456–469.e22. 10.1016/j.cell.2017.09.020

Kirst, C., Skriabine, S., Vieites-Prado, A., Topilko, T., Bertin, P., Gerschenfeld, G., Verny, F., Topilko, P., Michalski, N., Tessier-Lavigne, M., Renier, N., 2020. Mapping the Fine-Scale Organization and Plasticity of the Brain Vasculature. Cell 180, 780–795.e25. 10.1016/j.cell.2020.01.028

Klausberger, T., Somogyi, P., 2008. Neuronal Diversity and Temporal Dynamics: The Unity of Hippocampal Circuit Operations. Science 321, 53–57. 10.1126/science.1149381

Kleefstra, T., 2005. Disruption of the gene Euchromatin Histone Methyl Transferase1 (Eu-HMTase1) is associated with the 9q34 subtelomeric deletion syndrome. Journal of Medical Genetics 42, 299–306. 10.1136/jmg.2004.028464

Kleefstra, T., Brunner, H.G., Amiel, J., Oudakker, A.R., Nillesen, W.M., Magee, A., Geneviève, D., Cormier-Daire, V., Van Esch, H., Fryns, J.-P., Hamel, B.C.J., Sistermans, E.A., De Vries, B.B.A., Van Bokhoven, H., 2006. Loss-of-Function Mutations in Euchromatin Histone Methyl Transferase 1 (EHMT1) Cause the 9q34 Subtelomeric Deletion Syndrome. The American Journal of Human Genetics 79, 370–377. 10.1086/505693

Klein, S., Staring, M., Murphy, K., Viergever, M.A., Pluim, J., 2010. elastix: A Toolbox for Intensity-Based Medical Image Registration. IEEE Trans. Med. Imaging 29, 196–205. 10.1109/TMI.2009.2035616

Klingberg, A., Hasenberg, A., Ludwig-Portugall, I., Medyukhina, A., Männ, L., Brenzel, A., Engel, D.R., Figge, M.T., Kurts, C., Gunzer, M., 2017. Fully Automated Evaluation of Total Glomerular Number and Capillary Tuft Size in Nephritic Kidneys Using Lightsheet Microscopy. JASN 28, 452–459. 10.1681/ASN.2016020232

Kohlmeier, K.A., Reiner, P.B., 1999. Vasoactive Intestinal Polypeptide Excites Medial Pontine Reticular Formation Neurons in the Brainstem Rapid Eye Movement Sleep-Induction Zone. J. Neurosci. 19, 4073–4081. 10.1523/JNEUROSCI.19-10-04073.1999

Lee, B.R., Dalley, R., Miller, J.A., Chartrand, T., Close, J., Mann, R., Mukora, A., Ng, L., Alfiler, L., Baker, K., Bertagnolli, D., Brouner, K., Casper, T., Csajbok, E., Donadio, N., Driessens, S.L.W., Egdorf, T., Enstrom, R., Galakhova, A.A., Gary, A., Gelfand, E., Goldy, J., Hadley, K., Heistek, T.S., Hill, D., Hou, W.-H., Johansen, N., Jorstad, N., Kim, L., Kocsis, A.K., Kruse, L., Kunst, M., León, G., Long, B., Mallory, M., Maxwell, M., McGraw, M., McMillen, D., Melief, E.J., Molnar, G., Mortrud, M.T., Newman, D., Nyhus, J., Opitz-Araya, X., Ozsvár, A., Pham, T., Pom, A., Potekhina, L., Rajanbabu, R., Ruiz, A., Sunkin, S.M., Szöts, I., Taskin, N., Thyagarajan, B., Tieu, M., Trinh, J., Vargas, S., Vumbaco, D., Waleboer, F., Walling-Bell, S., Weed, N., Williams, G., Wilson, J., Yao, S., Zhou, T., Barzó, P., Bakken, T., Cobbs, C., Dee, N., Ellenbogen, R.G., Esposito, L., Ferreira, M., Gouwens, N.W., Grannan, B., Gwinn, R.P., Hauptman, J.S., Hodge, R., Jarsky, T., Keene, C.D., Ko, A.L., Korshoej, A.R., Levi, B.P., Meier, K., Ojemann, J.G., Patel, A., Ruzevick, J., Silbergeld, D.L., Smith, K., Sørensen, J.C., Waters, J., Zeng, H., Berg, J., Capogna, M., Goriounova, N.A., Kalmbach, B., De Kock, C.P.J., Mansvelder, H.D., Sorensen, S.A., Tamas, G., Lein, E.S., Ting, J.T., 2023. Signature morphoelectric properties of diverse GABAergic interneurons in the human neocortex. Science 382, eadf6484. 10.1126/science.adf6484

Li, J., Tanzillo, A.F., Pizzirusso, G., Caccavano, A., Chittajallu, R., Sohn, M., Abebe, D., Zhang, Y., Pelkey, K.A., Dale, R.K., McBain, C.J., Petros, T.J., 2026. Reducing methylation of histone 3.3 lysine 4 in the medial ganglionic eminence and hypothalamus recapitulates neurodevelopmental disorder phenotypes. Nat Commun 17, 2984. 10.1038/s41467-026-69248-9

Liao, D., Hessler, N.A., Malinow, R., 1995. Activation of postsynaptically silent synapses during pairing-induced LTP in CA1 region of hippocampal slice. Nature 375, 400–404. 10.1038/375400a0

Liu, Qiang, Bell, B.J., Kim, D.W., Lee, S.S., Keles, M.F., Liu, Qili, Blum, I.D., Wang, A.A., Blank, E.J., Xiong, J., Bedont, J.L., Chang, A.J., Issa, H., Cohen, J.Y., Blackshaw, S., Wu, M.N., 2023. A clock-dependent brake for rhythmic arousal in the dorsomedial hypothalamus. Nat Commun 14, 6381. 10.1038/s41467-023-41877-4

Loomba, S., Straehle, J., Gangadharan, V., Heike, N., Khalifa, A., Motta, A., Ju, N., Sievers, M., Gempt, J., Meyer, H.S., Helmstaedter, M., 2022. Connectomic comparison of mouse and human cortex. Science 377, eabo0924. 10.1126/science.abo0924

Marín, O., 2024. Parvalbumin interneuron deficits in schizophrenia. European Neuropsychopharmacology 82, 44–52. 10.1016/j.euroneuro.2024.02.010

Moissidis, M., Abbasova, L., Selten, M., Alis, R., Bernard, C., Domínguez-Canterla, Y., Oozeer, F., Qin, S., Kelly, A., Mòdol, L., Vasistha, N.A., Jones, B., Dhami, P., Khodosevich, K., Hamid, F., Lavender, P., Flames, N., Marín, O., 2025. A postnatal molecular switch drives activity-dependent maturation of parvalbumin interneurons. Cell. 10.1016/j.cell.2025.06.029

Mossink, B., Negwer, M., Schubert, D., Nadif Kasri, N., 2021. The emerging role of chromatin remodelers in neurodevelopmental disorders: a developmental perspective. Cell. Mol. Life Sci. 78, 2517–2563. 10.1007/s00018-020-03714-5

Murray, E., Cho, J.H., Goodwin, D., Ku, T., Swaney, J., Kim, S.-Y., Choi, H., Park, Y.-G., Park, J.-Y., Hubbert, A., McCue, M., Vassallo, S., Bakh, N., Frosch, M.P., Wedeen, V.J., Seung, H.S., Chung, K., 2015. Simple, Scalable Proteomic Imaging for High-Dimensional Profiling of Intact Systems. Cell 163, 1500–1514. 10.1016/j.cell.2015.11.025

Negwer, M., Bosch, B., Bormann, M., Hesen, R., Lütje, L., Aarts, L., Rossing, C., Nadif Kasri, N., Schubert, D., 2022. FriendlyClearMap: an optimized toolkit for mouse brain mapping and analysis. GigaScience 12, giad035. 10.1093/gigascience/giad035

Negwer, M., Piera, K., Hesen, R., Lütje, L., Aarts, L., Schubert, D., Nadif Kasri, N., 2020. EHMT1 regulates Parvalbumin-positive interneuron development and GABAergic input in sensory cortical areas. Brain Struct Funct 225, 2701–2716. 10.1007/s00429-020-02149-9

Neher, E., 2015. Merits and Limitations of Vesicle Pool Models in View of Heterogeneous Populations of Synaptic Vesicles. Neuron 87, 1131–1142. 10.1016/j.neuron.2015.08.038

Newmaster, K.T., Nolan, Z.T., Chon, U., Vanselow, D.J., Weit, A.R., Tabbaa, M., Hidema, S., Nishimori, K., Hammock, E.A.D., Kim, Y., 2020. Quantitative cellular-resolution map of the oxytocin receptor in postnatally developing mouse brains. Nat Commun 11, 1885. 10.1038/s41467-020-15659-1

Oh, S.W., Harris, J.A., Ng, L., Winslow, B., Cain, N., Mihalas, S., Wang, Q., Lau, C., Kuan, L., Henry, A.M., Mortrud, M.T., Ouellette, B., Nguyen, T.N., Sorensen, S.A., Slaughterbeck, C.R., Wakeman, W., Li, Y., Feng, D., Ho, A., Nicholas, E., Hirokawa, K.E., Bohn, P., Joines, K.M., Peng, H., Hawrylycz, M.J., Phillips, J.W., Hohmann, J.G., Wohnoutka, P., Gerfen, C.R., Koch, C., Bernard, A., Dang, C., Jones, A.R., Zeng, H., 2014. A mesoscale connectome of the mouse brain. Nature 508, 207–214. 10.1038/nature13186

Pan, C., Cai, R., Quacquarelli, F.P., Ghasemigharagoz, A., Lourbopoulos, A., Matryba, P., Plesnila, N., Dichgans, M., Hellal, F., Ertürk, A., 2016. Shrinkage-mediated imaging of entire organs and organisms using uDISCO. Nat Methods 13, 859–867. 10.1038/nmeth.3964

Perumal, M.B., Latimer, B., Xu, L., Stratton, P., Nair, S., Sah, P., 2021. Microcircuit mechanisms for the generation of sharp-wave ripples in the basolateral amygdala: A role for chandelier interneurons. Cell Reports 35, 109106. 10.1016/j.celrep.2021.109106

Piet, A., Ponvert, N., Ollerenshaw, D., Garrett, M., Groblewski, P.A., Olsen, S., Koch, C., Arkhipov, A., 2024. Behavioral strategy shapes activation of the Vip-Sst disinhibitory circuit in visual cortex. Neuron 112, 1876–1890.e4. 10.1016/j.neuron.2024.02.008

Preuss, F., Walker, F., Möck, M., Witte, M., Staiger, J.F., 2026. Inhibitory circuit motifs of cortical somatosensory layer 5 SST interneurons are uniform within layers but specific across layers. The Journal of Physiology 604, 479–502. 10.1113/JP288309

Prichard, J.R., Stoffel, R.T., Quimby, D.L., Obermeyer, W.H., Benca, R.M., Behan, M., 2002. Fos immunoreactivity in rat subcortical visual shell in response to illuminance changes. Neuroscience 114, 781–793. 10.1016/S0306-4522(02)00293-2

Prönneke, A., Scheuer, B., Wagener, R.J., Möck, M., Witte, M., Staiger, J.F., 2015. Characterizing VIP Neurons in the Barrel Cortex of VIPcre/tdTomato Mice Reveals Layer-Specific Differences. Cereb. Cortex 25, 4854–4868. 10.1093/cercor/bhv202

Rachel, J., Möck, M., Daigle, T.L., Tasic, B., Witte, M., Staiger, J.F., 2025. VIP-to-SST Cell Circuit Motif Shows Differential Short-Term Plasticity across Sensory Areas of Mouse Cortex. J. Neurosci. 45, e0949242025. 10.1523/JNEUROSCI.0949-24.2025

Reéb, Z., Magyar, D., Weisz, F., Fekete, Z., Müller, K., Vikór, A., Péterfi, Z., Andrási, T., Veres, J.M., Hájos, N., 2025. Morphological and electrophysiological diversity and connectivity of principal neurons in the lateral and basal nuclei of the mouse amygdala. Sci Rep 15, 33675. 10.1038/s41598-025-18411-1

Reh, R.K., Dias, B.G., Nelson, C.A., Kaufer, D., Werker, J.F., Kolb, B., Levine, J.D., Hensch, T.K., 2020. Critical period regulation across multiple timescales. Proc. Natl. Acad. Sci. U.S.A. 117, 23242–23251. 10.1073/pnas.1820836117

Ren, S., Zhang, C., Yue, F., Tang, J., Zhang, W., Zheng, Y., Fang, Y., Wang, N., Song, Z., Zhang, Z., Zhang, X., Qin, H., Wang, Y., Xia, J., Jiang, C., He, C., Luo, F., Hu, Z., 2024. A midbrain GABAergic circuit constrains wakefulness in a mouse model of stress. Nat Commun 15, 2722. 10.1038/s41467-024-46707-9

Renier, N., Adams, E.L., Kirst, C., Wu, Z., Azevedo, R., Kohl, J., Autry, A.E., Kadiri, L., Umadevi Venkataraju, K., Zhou, Y., Wang, V.X., Tang, C.Y., Olsen, O., Dulac, C., Osten, P., Tessier-Lavigne, M., 2016. Mapping of Brain Activity by Automated Volume Analysis of Immediate Early Genes. Cell 165, 1789–1802. 10.1016/j.cell.2016.05.007

Renier, N., Wu, Z., Simon, D.J., Yang, J., Ariel, P., Tessier-Lavigne, M., 2014. iDISCO: A Simple, Rapid Method to Immunolabel Large Tissue Samples for Volume Imaging. Cell 159, 896–910. 10.1016/j.cell.2014.10.010

Rhodes, C.T., Thompson, J.J., Mitra, A., Asokumar, D., Lee, D.R., Lee, D.J., Zhang, Y., Jason, E., Dale, R.K., Rocha, P.P., Petros, T.J., 2022. An epigenome atlas of neural progenitors within the embryonic mouse forebrain. Nat Commun 13, 4196. 10.1038/s41467-022-31793-4

Rots, D., Bouman, Arianne, Yamada, A., Levy, M., Dingemans, A.J.M., De Vries, B.B.A., Ruiterkamp-Versteeg, M., De Leeuw, N., Ockeloen, C.W., Pfundt, R., De Boer, E., Kummeling, J., Van Bon, B., Van Bokhoven, H., Kasri, N.N., Venselaar, H., Alders, M., Kerkhof, J., McConkey, H., Kuechler, A., Elffers, B., Van Beeck Calkoen, R., Hofman, S., Smith, A., Valenzuela, M.I., Srivastava, S., Frazier, Z., Maystadt, I., Piscopo, C., Merla, G., Balasubramanian, M., Santen, G.W.E., Metcalfe, K., Park, S.-M., Pasquier, L., Banka, S., Donnai, D., Weisberg, D., Strobl-Wildemann, G., Wagemans, A., Vreeburg, M., Baralle, D., Foulds, N., Scurr, I., Brunetti-Pierri, N., Van Hagen, J.M., Bijlsma, E.K., Hakonen, A.H., Courage, C., Genevieve, D., Pinson, L., Forzano, F., Deshpande, C., Kluskens, M.L., Welling, L., Plomp, A.S., Vanhoutte, E.K., Kalsner, L., Hol, J.A., Putoux, A., Lazier, J., Vasudevan, P., Ames, E., O’Shea, J., Lederer, D., Fleischer, J., O’Connor, M., Pauly, M., Vasileiou, G., Reis, A., Kiraly-Borri, C., Bouman, Arjan, Barnett, C., Nezarati, M., Borch, L., Beunders, G., Özcan, K., Miot, S., Volker-Touw, C.M.L., Van Gassen, K.L.I., Cappuccio, G., Janssens, K., Mor, N., Shomer, I., Dominissini, D., Tedder, M.L., Muir, A.M., Sadikovic, B., Brunner, H.G., Vissers, L.E.L.M., Shinkai, Y., Kleefstra, T., 2024. Comprehensive EHMT1 variants analysis broadens genotype-phenotype associations and molecular mechanisms in Kleefstra syndrome. The American Journal of Human Genetics 111, 1605–1625. 10.1016/j.ajhg.2024.06.008

Rudy, B., Fishell, G., Lee, S., Hjerling-Leffler, J., 2011. Three groups of interneurons account for nearly 100% of neocortical GABAergic neurons. Developmental Neurobiology 71, 45–61. 10.1002/dneu.20853

Schaefer, A., Sampath, S.C., Intrator, A., Min, A., Gertler, T.S., Surmeier, D.J., Tarakhovsky, A., Greengard, P., 2009. Control of Cognition and Adaptive Behavior by the GLP/G9a Epigenetic Suppressor Complex. Neuron 64, 678–691. 10.1016/j.neuron.2009.11.019

Schut, E.H.S., Alonso, A., Smits, S., Khamassi, M., Samanta, A., Negwer, M., Kasri, N.N., Navarro Lobato, I., Genzel, L., 2020. The Object Space Task reveals increased expression of cumulative memory in a mouse model of Kleefstra syndrome. Neurobiology of Learning and Memory 173, 107265. 10.1016/j.nlm.2020.107265

Segal, M., 2010. Dendritic spines, synaptic plasticity and neuronal survival: activity shapes dendritic spines to enhance neuronal viability. Eur J of Neuroscience 31, 2178–2184. 10.1111/j.1460-9568.2010.07270.x

Spampanato, J., Sullivan, R.K.P., Perumal, M.B., Sah, P., 2016. Development and physiology of GABAergic feedback excitation in parvalbumin expressing interneurons of the mouse basolateral amygdala. Physiol Rep 4, e12664. 10.14814/phy2.12664

Sun, Y., Qian, L., Xu, L., Hunt, S., Sah, P., 2023. Somatostatin neurons in the central amygdala mediate anxiety by disinhibition of the central sublenticular extended amygdala. Mol Psychiatry 28, 4163–4174. 10.1038/s41380-020-00894-1

Szkudlarek, H.J., Herdzina, O., Lewandowski, M.H., 2008. Ultra-slow oscillatory neuronal activity in the rat olivary pretectal nucleus: comparison with oscillations within the intergeniculate leaflet. Eur J of Neuroscience 27, 2657–2664. 10.1111/j.1460-9568.2008.06225.x

Szkudlarek, H.J., Orlowska, P., Lewandowski, M.H., 2012. Light-Induced Responses of Slow Oscillatory Neurons of the Rat Olivary Pretectal Nucleus. PLoS ONE 7, e33083. 10.1371/journal.pone.0033083

Tachibana, M., Matsumura, Y., Fukuda, M., Kimura, H., Shinkai, Y., 2008. G9a/GLP complexes independently mediate H3K9 and DNA methylation to silence transcription. EMBO J 27, 2681–2690. 10.1038/emboj.2008.192

Tachibana, M., Sugimoto, K., Nozaki, M., Ueda, J., Ohta, T., Ohki, M., Fukuda, M., Takeda, N., Niida, H., Kato, H., Shinkai, Y., 2002. G9a histone methyltransferase plays a dominant role in euchromatic histone H3 lysine 9 methylation and is essential for early embryogenesis. Genes Dev. 16, 1779–1791. 10.1101/gad.989402

Tachibana, M., Ueda, J., Fukuda, M., Takeda, N., Ohta, T., Iwanari, H., Sakihama, T., Kodama, T., Hamakubo, T., Shinkai, Y., 2005. Histone methyltransferases G9a and GLP form heteromeric complexes and are both crucial for methylation of euchromatin at H3-K9. Genes Dev. 19, 815–826. 10.1101/gad.1284005

Tang, X., Jaenisch, R., Sur, M., 2021. The role of GABAergic signalling in neurodevelopmental disorders. Nat Rev Neurosci 22, 290–307. 10.1038/s41583-021-00443-x

Todd, W.D., Venner, A., Anaclet, C., Broadhurst, R.Y., De Luca, R., Bandaru, S.S., Issokson, L., Hablitz, L.M., Cravetchi, O., Arrigoni, E., Campbell, J.N., Allen, C.N., Olson, D.P., Fuller, P.M., 2020. Suprachiasmatic VIP neurons are required for normal circadian rhythmicity and comprised of molecularly distinct subpopulations. Nat Commun 11, 4410. 10.1038/s41467-020-17197-2

Tremblay, R., Lee, S., Rudy, B., 2016. GABAergic Interneurons in the Neocortex: From Cellular Properties to Circuits. Neuron 91, 260–292. 10.1016/j.neuron.2016.06.033

Trouche, S., Sasaki, J.M., Tu, T., Reijmers, L.G., 2013. Fear Extinction Causes Target-Specific Remodeling of Perisomatic Inhibitory Synapses. Neuron 80, 1054–1065. 10.1016/j.neuron.2013.07.047

Vermeulen, K., De Boer, A., Janzing, J.G.E., Koolen, D.A., Ockeloen, C.W., Willemsen, M.H., Verhoef, F.M., Van Deurzen, P.A.M., Van Dongen, L., Van Bokhoven, H., Egger, J.I.M., Staal, W.G., Kleefstra, T., 2017a. Adaptive and maladaptive functioning in Kleefstra syndrome compared to other rare genetic disorders with intellectual disabilities. American J of Med Genetics Pt A 173, 1821–1830. 10.1002/ajmg.a.38280

Vermeulen, K., Staal, W.G., Janzing, J.G., Van Bokhoven, H., Egger, J.I.M., Kleefstra, T., 2017b. Sleep Disturbance as a Precursor of Severe Regression in Kleefstra Syndrome Suggests a Need for Firm and Rapid Pharmacological Treatment. Clin Neuropharm 40, 185–188. 10.1097/WNF.0000000000000226

Wang, Q., Bhandiwad, A., Gouwens, N.W., Yao, S., Wang, Y., Kuang, X., Li, A., Li, X., Dalley, R., Kuo, H.-C., Lesnar, P., Xu, W., Mallory, M., Li, Y., El-Hifnawi, L., Ahmadinia, L., Ouellette, B., Kruse, L., Ng, L., Gong, H., Luo, Q., Kunst, M., Van Velthoven, C.T.J., Yao, Z., Sorensen, S.A., Zeng, H., 2026. Genoarchitecture and input–output organization of the mouse basal ganglia and thalamic parafascicular nucleus. Nat Neurosci 29, 1248–1264. 10.1038/s41593-026-02253-9

Wang, Q., Ding, S.-L., Li, Y., Royall, J., Feng, D., Lesnar, P., Graddis, N., Naeemi, M., Facer, B., Ho, A., Dolbeare, T., Blanchard, B., Dee, N., Wakeman, W., Hirokawa, K.E., Szafer, A., Sunkin, S.M., Oh, S.W., Bernard, A., Phillips, J.W., Hawrylycz, M., Koch, C., Zeng, H., Harris, J.A., Ng, L., 2020. The Allen Mouse Brain Common Coordinate Framework: A 3D Reference Atlas. Cell 181, 936–953.e20. 10.1016/j.cell.2020.04.007

Willemsen, M.H., Vulto-van Silfhout, A.T., Nillesen, W.M., Wissink-Lindhout, W.M., Van Bokhoven, H., Philip, N., Berry-Kravis, E.M., Kini, U., Van Ravenswaaij-Arts, C.M.A., Delle Chiaie, B., Innes, A.M.M., Houge, G., Kosonen, T., Cremer, K., Fannemel, M., Stray-Pedersen, A., Reardon, W., Ignatius, J., Lachlan, K., Mircher, C., Helderman Van Den Enden, P.T.J.M., Mastebroek, M., Cohn-Hokke, P.E., Yntema, H.G., Drunat, S., Kleefstra, T., 2011. Update on Kleefstra Syndrome. Mol Syndromol 2, 202–212. 10.1159/000335648

Williams, L.E., Holtmaat, A., 2019. Higher-Order Thalamocortical Inputs Gate Synaptic Long-Term Potentiation via Disinhibition. Neuron 101, 91–102.e4. 10.1016/j.neuron.2018.10.049

Wong, F.K., Bercsenyi, K., Sreenivasan, V., Portalés, A., Fernández-Otero, M., Marín, O., 2018. Pyramidal cell regulation of interneuron survival sculpts cortical networks. Nature 557, 668–673. 10.1038/s41586-018-0139-6

Wu, S.J., Sevier, E., Dwivedi, D., Saldi, G.-A., Hairston, A., Yu, S., Abbott, L., Choi, D.H., Sherer, M., Qiu, Y., Shinde, A., Lenahan, M., Rizzo, D., Xu, Q., Barrera, I., Kumar, V., Marrero, G., Prönneke, A., Huang, S., Kullander, K., Stafford, D.A., Macosko, E., Chen, F., Rudy, B., Fishell, G., 2023. Cortical somatostatin interneuron subtypes form cell-type-specific circuits. Neuron 111, 2675–2692.e9. 10.1016/j.neuron.2023.05.032

Yamada, A., Hirasawa, T., Nishimura, K., Shimura, C., Kogo, N., Fukuda, K., Kato, M., Yokomori, M., Hayashi, T., Umeda, M., Yoshimura, M., Iwakura, Y., Nikaido, I., Itohara, S., Shinkai, Y., 2021. Derepression of inflammation-related genes link to microglia activation and neural maturation defect in a mouse model of Kleefstra syndrome. iScience 24, 102741. 10.1016/j.isci.2021.102741

Yao, Z., Liu, H., Xie, F., Fischer, S., Adkins, R.S., Aldridge, A.I., Ament, S.A., Bartlett, A., Behrens, M.M., Van Den Berge, K., Bertagnolli, D., De Bézieux, H.R., Biancalani, T., Booeshaghi, A.S., Bravo, H.C., Casper, T., Colantuoni, C., Crabtree, J., Creasy, H., Crichton, K., Crow, M., Dee, N., Dougherty, E.L., Doyle, W.I., Dudoit, S., Fang, R., Felix, V., Fong, O., Giglio, M., Goldy, J., Hawrylycz, M., Herb, B.R., Hertzano, R., Hou, X., Hu, Q., Kancherla, J., Kroll, M., Lathia, K., Li, Y.E., Lucero, J.D., Luo, C., Mahurkar, A., McMillen, D., Nadaf, N.M., Nery, J.R., Nguyen, T.N., Niu, S.-Y., Ntranos, V., Orvis, J., Osteen, J.K., Pham, T., Pinto-Duarte, A., Poirion, O., Preissl, S., Purdom, E., Rimorin, C., Risso, D., Rivkin, A.C., Smith, K., Street, K., Sulc, J., Svensson, V., Tieu, M., Torkelson, A., Tung, H., Vaishnav, E.D., Vanderburg, C.R., Van Velthoven, C., Wang, X., White, O.R., Huang, Z.J., Kharchenko, P.V., Pachter, L., Ngai, J., Regev, A., Tasic, B., Welch, J.D., Gillis, J., Macosko, E.Z., Ren, B., Ecker, J.R., Zeng, H., Mukamel, E.A., 2021. A transcriptomic and epigenomic cell atlas of the mouse primary motor cortex. Nature 598, 103–110. 10.1038/s41586-021-03500-8

Yau, C.N., Hung, J.T.S., Campbell, R.A.A., Wong, T.C.Y., Huang, B., Wong, B.T.Y., Chow, N.K.N., Zhang, L., Tsoi, E.P.L., Tan, Y., Li, J.J.X., Wing, Y.K., Lai, H.M., 2024. INSIHGT: an accessible multi-scale, multi-modal 3D spatial biology platform. Nat Commun 15, 10888. 10.1038/s41467-024-55248-0

